# *ADCY3* Ser107Pro links difficulty awakening in the morning to adiposity through circadian regulation of adipose thermogenesis

**DOI:** 10.1101/2025.07.28.667339

**Authors:** Cynthia Tchio, Matthew Maher, Christopher Moth, Jens Meiler, Jacqueline Lane, Herman Taylor, Jonathan Williams, Richa Saxena

## Abstract

Modern lifestyles often disturb circadian rhythms, yet the genetic circuits that convert this stress into metabolic dysfunction remain poorly defined. Here, we identify a missense variant in *ADCY3* (rs11676272; Ser107Pro) as a pleiotropic regulator of circadian preference and adiposity. Using genome-wide pleiotropy analysis in ∼480,000 UK Biobank participants, we show that the G risk allele (Pro107) increases morningness, BMI, and fat mass in European (N=451,324) and African (N=8,738) ancestry groups, with behavioral amplification by morning difficulty awakening observed in Europeans, power limited interaction modeling in other populations. Structural modeling and transcriptomic analysis suggest this allele destabilizes ADCY3 and alters adipose-specific splicing and expression. In mice, *Adcy3* is rhythmically expressed in adipose tissue, with the conserved Pro107 site showing BMAL1 binding and cold-inducible activation. Human adipose *ADCY3* expression also increases after weight loss. Together, these findings reveal a genotype-dependent, behaviorally modifiable axis linking difficulty awakening to adipose thermogenesis and obesity risk.

## Graphical Abstract Summary

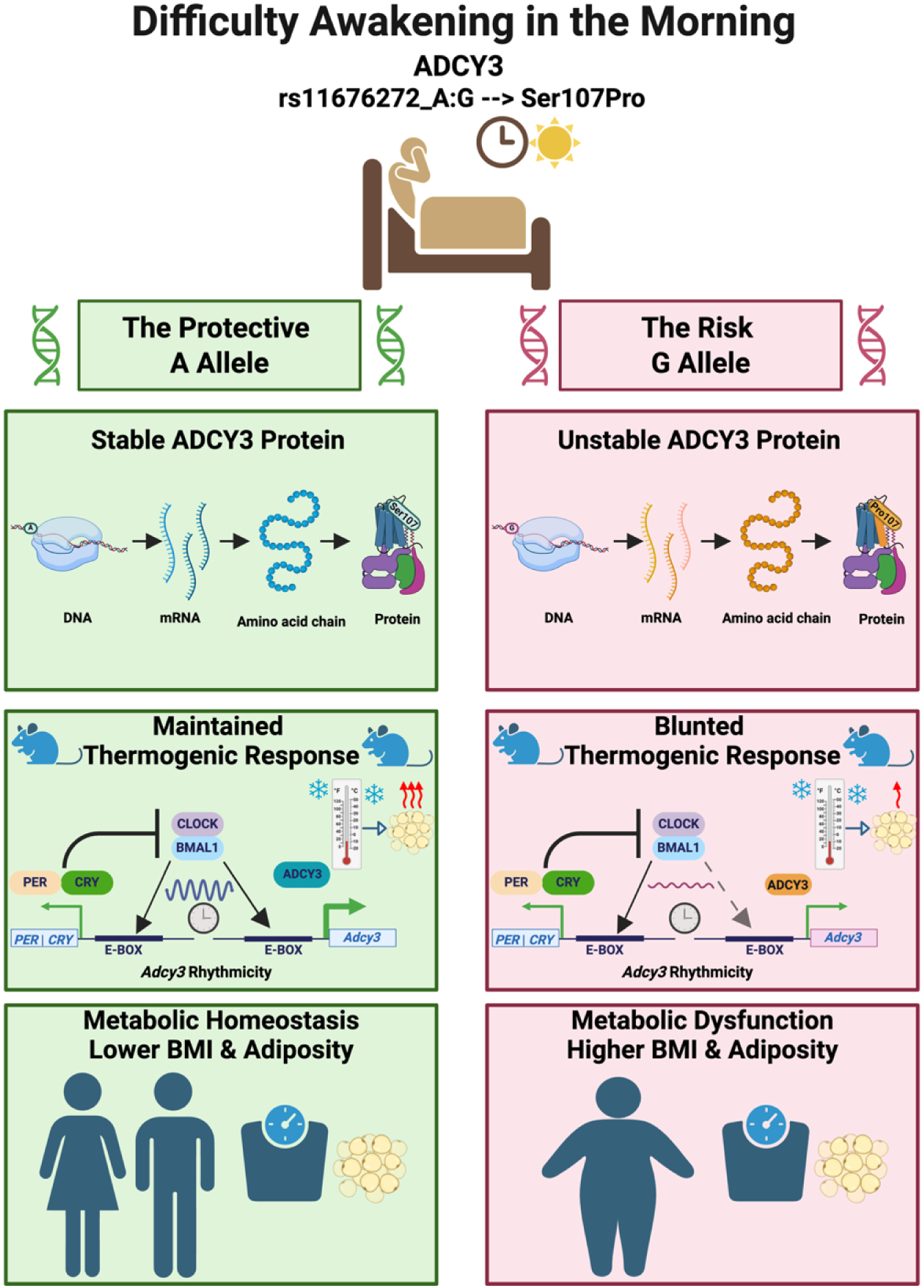
Difficulty awakening in the morning is associated with increased adiposity, and we show this susceptibility is modulated by a common variant in ADCY3 (rs11676272). The risk G allele encodes a destabilizing Ser107Pro missense variant, predicted to impair protein function in adipose tissue, where ADCY3 is rhythmically expressed under the control of BMAL1. This may blunt the thermogenic response and promote metabolic dysfunction. In contrast, the protective A allele supports the stable expression of ADCY3 protein, preserves rhythmicity, and promotes normal thermogenesis. Together, our results reveal a mechanistic model linking genetic variation, circadian regulation, and behavioral strain to obesity risk.

## INTRODUCTION

The circadian clock governs a wide range of physiological processes, including sleep-wake cycles, metabolism, and energy homeostasis ^1,2^. Chronotype reflects an individual’s behavioral preference for morning or evening activity, summarizing the phase of the circadian system as shaped by both environmental and genetic factors ^3^. Genome-wide association studies (GWAS) have identified over 300 loci for chronotype; many of which colocalize with loci for BMI and fat mass, pointing to molecular crosstalk between circadian and metabolic regulation ^1,3–7^.

A related but less-studied trait is self-reported difficulty waking in the morning, which reflects a person’s ability to arouse from sleep on schedule. Although genetically correlated with chronotype ^6^ , recent work suggests that this trait captures a distinct dimension of circadian strain, potentially linked to misalignment and impaired arousal regulation ^8^. Understanding how genetic variation modifies this trait could reveal clock-controlled metabolic pathways that are particularly sensitive to behavioral disruption.

The canonical clock genes offer proof of principle, where the deletion of Clock or Bmal1 (Arntl) in mice leads to obesity and glucose intolerance ^9,10^, and clock output in brown and white adipose tissue gates lipid mobilization and thermogenesis ^11^. However, for the vast majority of human GWAS signals, the causal alleles, effector genes, and tissue-specific mechanisms remain unresolved. Notably, no prior study has identified a single coding variant that links a circadian behavioral trait and an adipose-centric metabolic phenotype through functional validation in relevant tissues.

Here, we use a genome-wide pleiotropy approach to bridge this gap. By jointly analyzing GWAS summary statistics for morning chronotype, difficulty waking in the morning, and BMI in ∼480,000 UK Biobank participants of European, African, and South Asian ancestry, we identify a shared missense signal at ADCY3 (rs11676272; Ser107Pro). We show that the BMI-raising G allele (Pro107) has a stronger effect in individuals who report difficulty waking, suggesting a gene-by-behavior interaction. To dissect the mechanism, we integrate in silico protein modeling, human eQTL/splice-QTL analysis, and mouse adipose physiology.

In humans, the G allele destabilizes ADCY3 protein and alters its splicing and expression in adipose tissue. In mice, *Adcy3* is rhythmically expressed in adipose, and its conserved Pro107 site is a BMAL1 binding target, with expression increasing during cold-induced thermogenesis. Furthermore, human adipose RNA-seq from weight-loss interventions shows that ADCY3 expression increases in metabolically improved states, reinforcing its role in energy balance.

Together, these findings position ADCY3 Ser107Pro as a molecular link between a daily arousal trait and adipose thermogenesis, illustrating how common genetic variation can amplify the metabolic cost of modern sleep behavior. We propose a unified model in which genetic variation in ADCY3 mediates the metabolic consequences of lifestyle and circadian strain, offering a functional framework that connects genetics, environment, and tissue-specific regulation to obesity risk.

## RESULTS

### Genome-wide pleiotropy analysis identifies shared loci between circadian preference traits and BMI

To identify genetic signals influencing both circadian rhythms and BMI, we conducted a genome-wide pleiotropy analysis using the PLACO composite-null test ^12^ and HyPrColoc colocalization ^13^ starting from GWAS summary statistics for morningness chronotype ^4^ (n = 449,734); ease of waking up in the morning (n = 451,872; both UKB Europeans), and BMI (n = 694,649; UKB and GIANT consortium^14^ ; **Table S1**). We identified 10 pleiotropic loci for morningness chronotype (Chrono) and BMI (**Figure 1A, Table S2**) and 5 loci for ease of waking up in the morning (Getup) and BMI (**Figure 1D, Table S3**). Four loci (in or near *ADCY3*, *CELF1*, *TFAP2B*, and *FTO*) were common to both analyses. Regional colocalization analysis using a different approach (HyPrColoc ^13^) confirmed that the same SNPs underlie associations across all three traits, with evidence of shared regulation across all three traits: morningness (Chrono), difficulty waking (Getup), and BMI (posterior probability, PP > 0.80, **Table 1**). Notably, pleiotropic variants associated with higher BMI are associated with either evening chronotype (*ADCY3*) or morning chronotype (*TFAP2B*, *CELF1, and FTO*). Phenome-wide association analysis (PheWAS) revealed that all four pleiotropic loci are also significantly associated with fat mass, highlighting their role in adiposity regulation (**Table S4**). Furthermore, *in silico* assessment revealed that the lead SNPs reside in active regulatory regions, demonstrated by tissue-specific chromatin interactions (**Figure S2**) and significant eQTL effects (**Table S5**).

**Figure 1:**
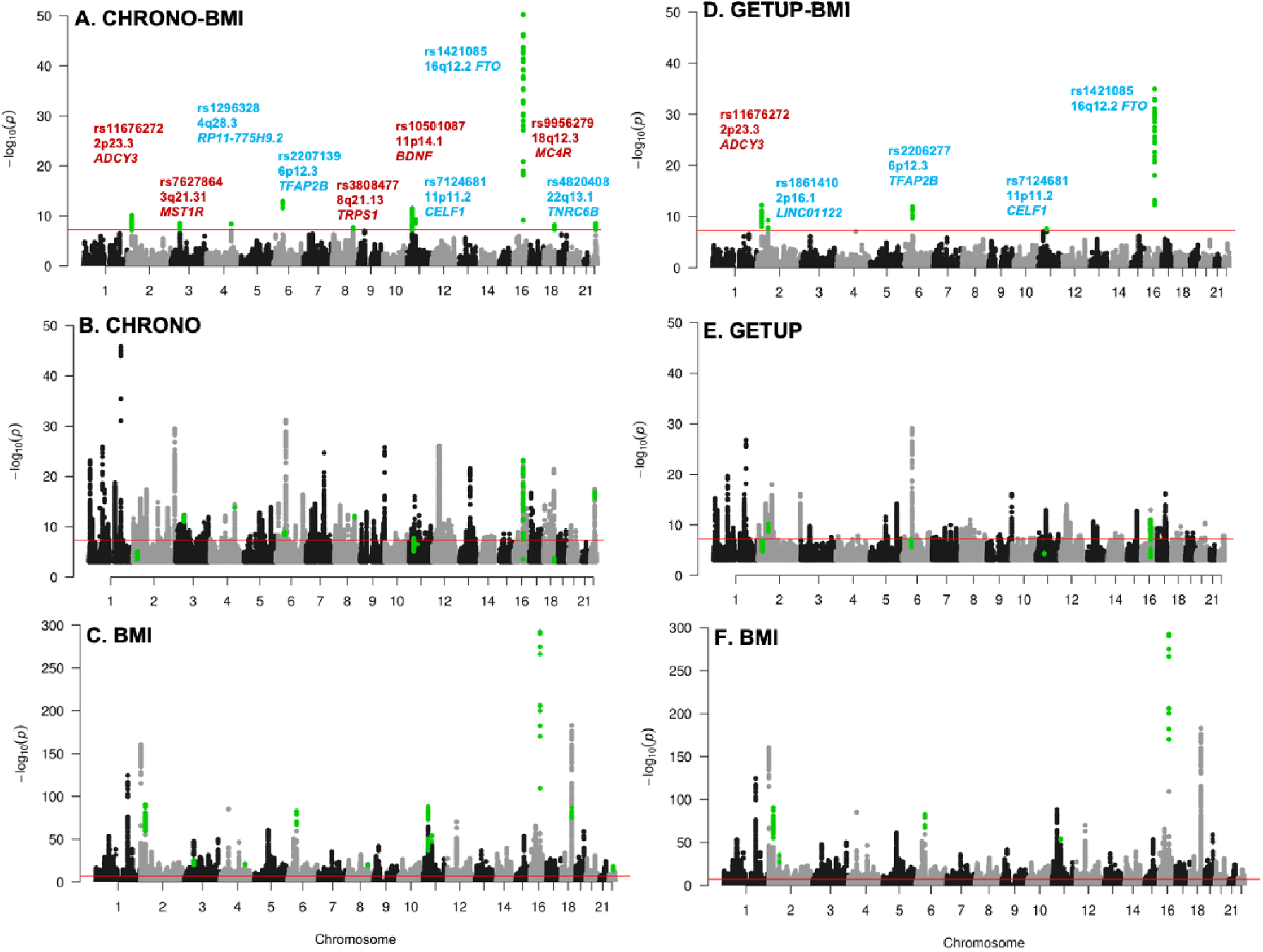
Genome-wide analysis identifies shared genetic loci between circadian traits and BMI. Manhattan plots showing genome-wide pleiotropy for (**A**) morningness chronotype × BMI and (**D**) ease of waking up in the morning × BMI. Single-trait GWAS plots shown for (**B**) chronotype, (**E**) ease of waking up in the morning, and (**C, F**) BMI. The red line marks the genome-wide significance threshold (p < 5×10⁻ ). In (**A**) and (**D**), pleiotropic loci are annotated with the nearest gene; red highlights loci with opposite directions of effect across traits, while blue indicates shared directionality.

**Table 1:** Colocalization identifies shared candidate causal loci for morning circadian preference traits and BMI.

| Gene | Traits | Posterior P | Regional P | Candidate SNP | Annotation | P by SNP |
| --- | --- | --- | --- | --- | --- | --- |
| *ADCY3* | Chrono, Getup, BMI | 0.843 | 0.924 | rs11676272 | Missense | 1.000 |
| *CELF1* | Chrono, Getup, BMI | 0.615 | 0.662 | rs7124681 | Intronic | 1.000 |
| *FTO* | Chrono, Getup, BMI | 0.972 | 1.000 | rs1558902 | Intronic | 0.917 |
| *TFAP2B* | Chrono, Getup, BMI | 0.991 | 0.999 | rs2206277 | Intronic | 0.828 |

Colocalization analyses were conducted using HyPrColoc (Hypothesis Prioritization in Multi-Trait Colocalization) and GWAS summary statistics (Chrono: Morningness Chronotype; Getup: Ease of waking up in the morning; BMI: Body Mass Index). 'Posterior P' (posterior probability of colocalization) assumes that a locus contains 1 causal variant, whereas 'Regional P' assumes that a locus contains >1 causal variant. 'P by SNP' is the % contribution of the candidate (causal) variant to the posterior p.

### A missense variant in *ADCY3* is the top candidate for shared causality

Of the four pleiotropic loci, we focus on rs11676272 in *ADCY3*, a missense variant (Ser107Pro), because it showed a strong colocalization across all three traits, with a PP of 0.84, with the SNP contributing to 100% of the PP (**Table 1**). Specifically, the G allele, which encodes for Proline-107, associates with eveningness (Chrono β = -0.0118 ± 0.0026), more difficulty waking up in the morning (Getup β = -0.0084 ± 0.00157), and higher BMI (β = 0.0328 ± 0.002); the regional plots for all three traits are shown in **Figure 2**. This SNP was prioritized not only for its robust statistical signal but also because its dual potential to alter ADCY3 protein structure while also regulating gene expression provided a unique opportunity to investigate multi-level gene function. Population frequency analysis revealed that rs11676272 varies across ancestries (33-85% in GnomAD), with the ancestral G allele being most prevalent in African populations (**Figure S10A**). The A allele frequency varies across populations and appears to have undergone recent positive selection (**Figure S10B**) based on Tajima’s D statistics (AFR: –1.33; EAS: –1.44; EUR: –0.71), consistent with adaptation to cold environments.

**Figure 2:**
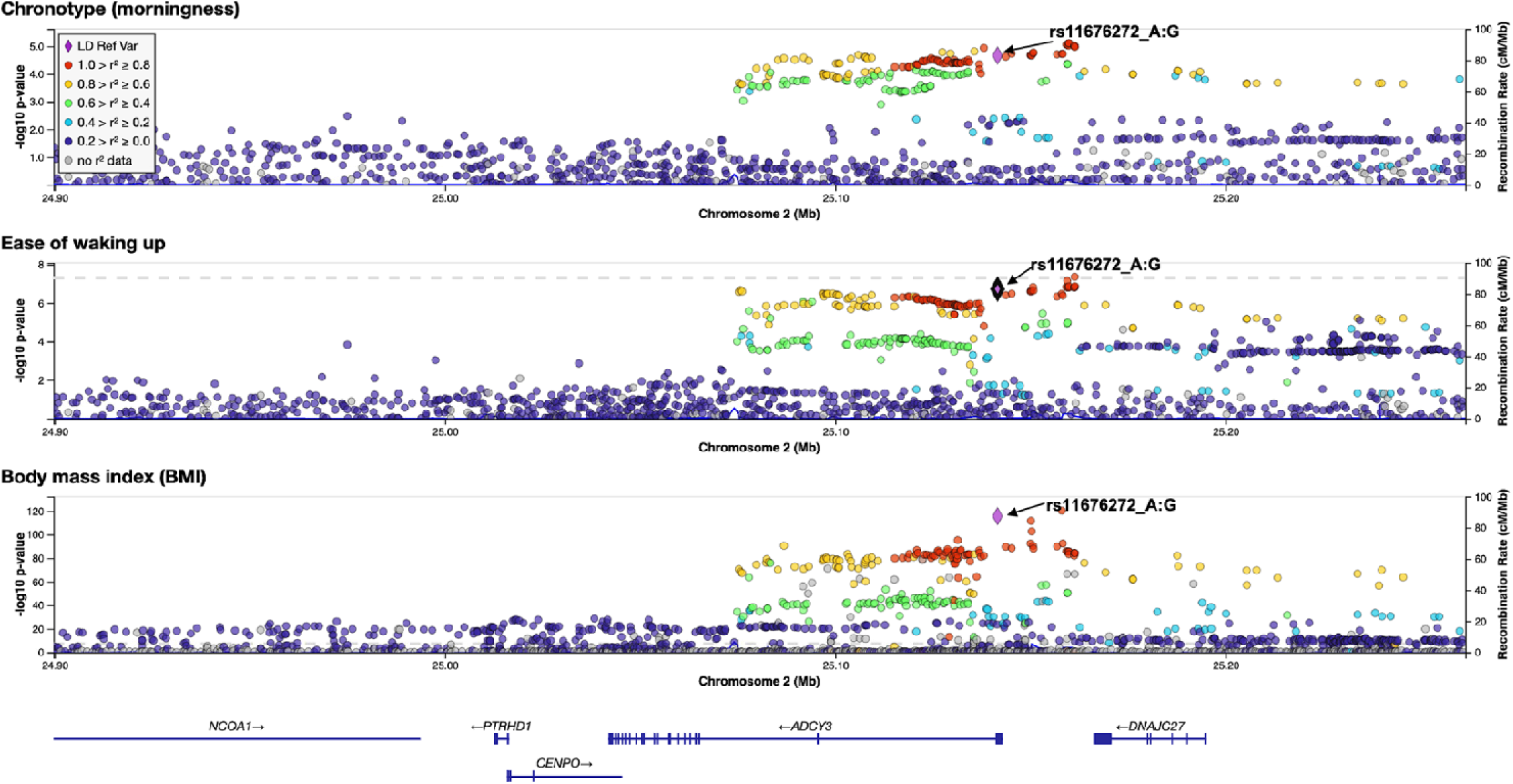
Colocalization analysis supports rs11676272 as a shared causal variant for circadian traits and BMI at the *ADCY3* locus. Regional plots for (Top) chronotype, (Middle) ease of waking, and (Bottom) BMI. The lead pleiotropic missense SNP rs11676272 is marked with a purple diamond. Colors reflect linkage disequilibrium (r²) with rs11676272, based on the 1000 Genomes EUR reference panel. Colocalization analysis (HyPrColoc) supports a shared causal variant across all traits (PP = 0.84; regional PP = 0.92; SNP PP = 1.00). Genomic region spans chr2:24,992,038– 25,193,106 (GRCh37).

### The ADCY3 risk allele’s effect on adiposity is sexually dimorphic and is amplified by difficulty awakening in the morning

To dissect the lead pleiotropic variant, rs11676272, we performed linear mixed models using lme4^15^ in the UK-Biobank cohort. The G risk allele, previously linked to eveningness, showed a clear dose-response with higher BMI, fat mass, and body-fat percentage (**Figure 3A-C**, **Figure S3A-B**). The effect was largest in Europeans (AG vs GG β = –0.19 kg m⁻²; AA vs GG β = –0.32 kg m⁻²; P < 2 × 10⁻¹ ) and directionally concordant in Africans (AG vs GG β = –0.38 kg m⁻²; P = 0.009), but imprecise in South Asians (**Table S8**). Thus, each A allele lowers BMI relative to the G risk allele. A clear sex-dimorphic pattern emerged: the protective A allele produced a larger reduction in females (AA vs GG β = –0.38 kg m⁻²) than in males (β = –0.24 kg m⁻²; P for gene-by-sex interaction = 2.4 × 10⁻¹ ; **Figure 3B**). The same female-greater trend was observed in Africans (**Figure 3D**).

**Figure 3:**
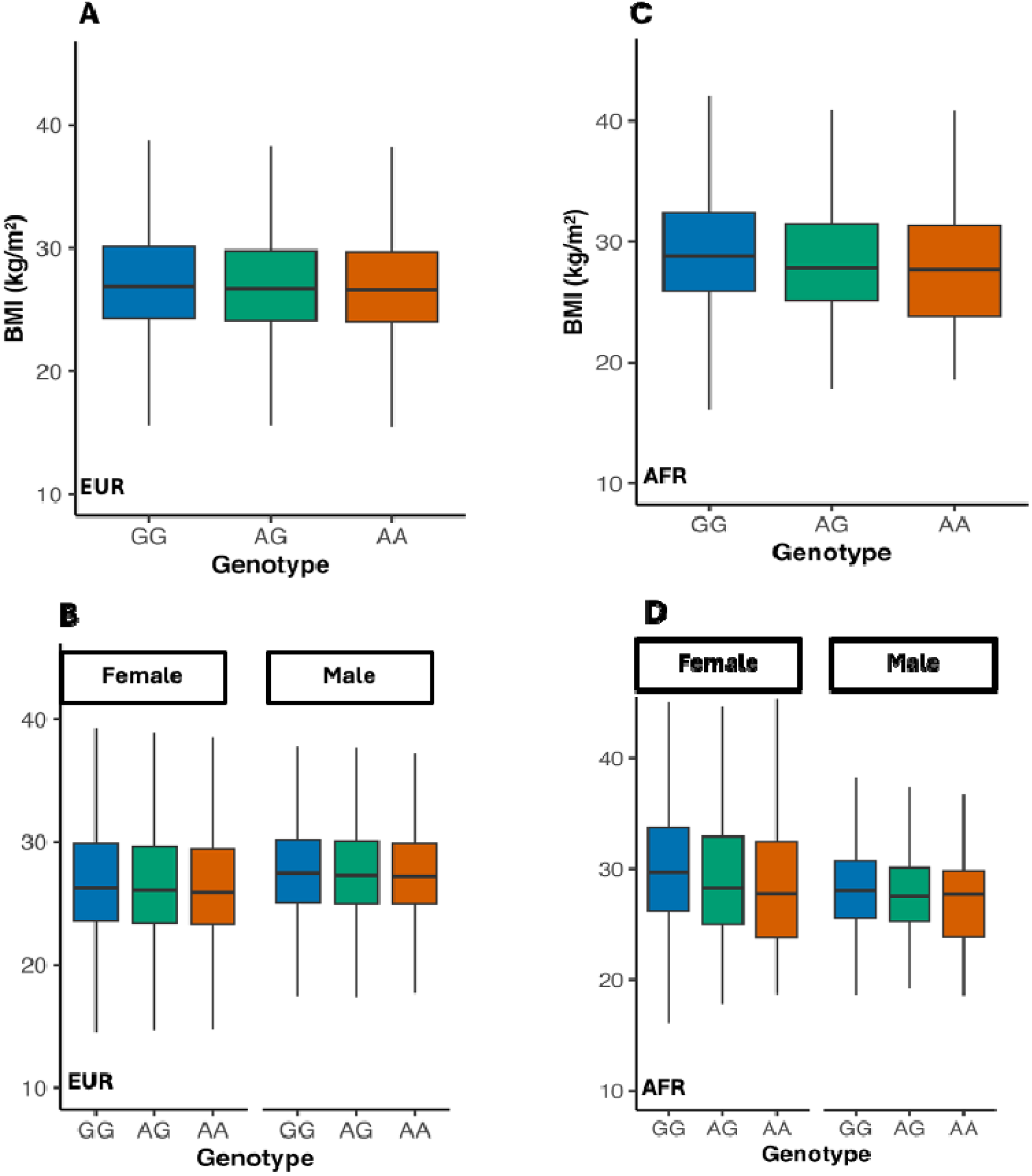
The *ADCY3* risk allele G associates with higher BMI in both Europeans and Africans, with stronger effect in women. (**A**) In Europeans (N = 451,324), BMI decreases with each additional A allele (AA vs GG = –0.32 kg/m², P < 2×10⁻¹ ). (**B**) Sex-stratified analysis shows a stronger protective effect in females (AA vs GG = –0.38) than males (–0.24), with significant gene-by-sex interaction (P = 2.4×10⁻¹ ). (**C**) In Africans (N = 8,738), AG carriers have lower BMI than GG (β = –0.38, P = 0.009); AA is underpowered. (**D**) Sex-stratified analysis in Africans shows a similar female-greater trend. Box plots: center = median; box = IQR; whiskers = 1.5×IQR. Colors: GG (blue), AG (green), AA (orange).

We next asked whether self-reported “difficulty awakening in the morning” modifies this genetic effect. In Europeans, we observed a significant interaction for both BMI (AG × GET_UP β = –0.044, P = 0.049) and fat mass (AA × GET_UP β = –0.153, P = 0.002) (**Figures 4A, 4B**). The positive association between increased difficulty awakening in the morning and higher adiposity was most pronounced in G-allele carriers, whereas A-allele homozygotes were largely protected. No significant interaction was observed for chronotype preference itself (**Figure S4, Table S8**), nor for night shift work (n = 42,486; all P > 0.1; **Table S8**), underscoring the specificity of the morning-waking trait.

**Figure 4:**
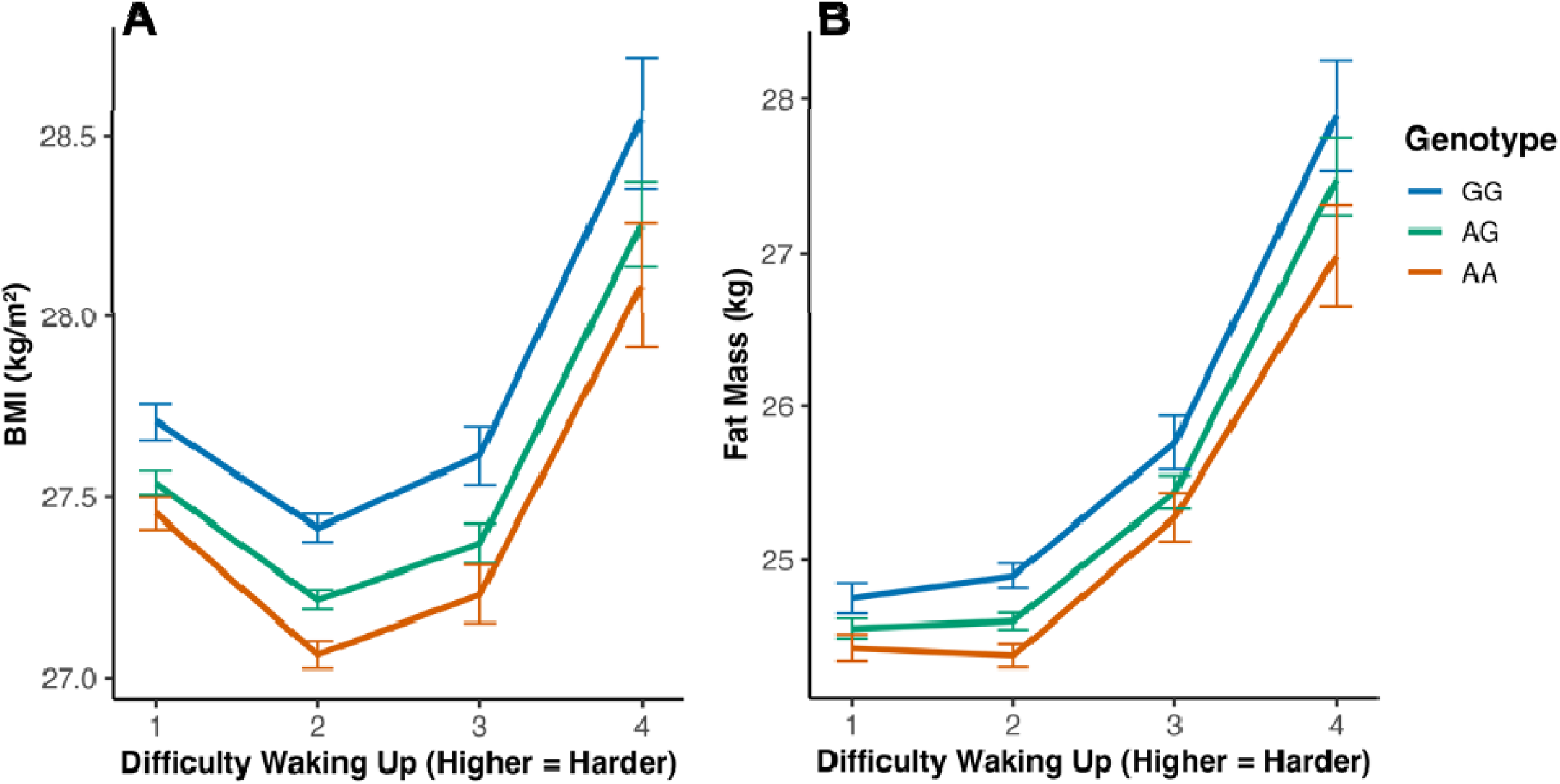
The obesogenic effect of the *ADCY3* risk allele is amplified by difficulty waking up in the morning. Interaction analysis between rs11676272 genotype and waking difficulty in UK Biobank Europeans (N = 451,324). (**A**) Effect on BMI. (**B**) Effect on fat mass. Genotype-stratified slopes show that the association between waking difficulty and adiposity is strongest in GG carriers and blunted in AA homozygotes. Models adjusted for age, sex, PC1–10, and kinship. Error bars = 95% CI.

For African and South Asian participants, the allele showed the same protective direction in heterozygotes (AG vs GG, β = -0.38 kg m⁻², P = 0.009 in Africans; β = -0.07 kg m⁻² , P = 0.568 in South Asians) (**Table S8**). Sample-size differences explain the wider confidence intervals in these groups: Europeans contribute > 120,000 AA homozygotes, Africans <150, and South Asians ∼2,000 (**Figure S5**).

Together, these data demonstrate that rs11676272 influences adiposity in a sex-dimorphic manner and that, in Europeans, its obesogenic effect is exacerbated by difficulty waking up in the morning, a behaviorally tractable facet of circadian physiology. **The rs11676272 variant has multi-level functional consequences.**

To elucidate the molecular mechanisms of rs11676272, we first modeled the structural consequences of the Ser107Pro missense mutation. Protein modeling predicted that the substitution of proline for serine disrupts key hydrogen bonds and significantly destabilizes the ADCY3 protein structure (**Figure 5A-C**). In addition to the protein-altering effect, we found that rs11676272 is a significant expression and splicing quantitative trait locus (eQTL and sQTL) for *ADCY3* in subcutaneous adipose tissue (**Figure 5D** and **Figure S7A**). The risk allele G was paradoxically associated with higher *ADCY3* expression, potentially reflecting a compensatory transcriptional response to impaired protein stability or altered feedback regulation. Mendelian randomization analysis supported a causal relationship between this altered *ADCY3* expression and the observed traits (**Figure 5E**). The variant’s location within an active regulatory element in adipose tissue, confirmed by histone markers H3K4me1 and H3K4me3 (**Figure S6**), and binding site to multiple transcription factors, including RNA Polymerase II POLR2A (**Table S7**), provides a strong mechanistic basis for its regulatory effects.

**Figure 5:**
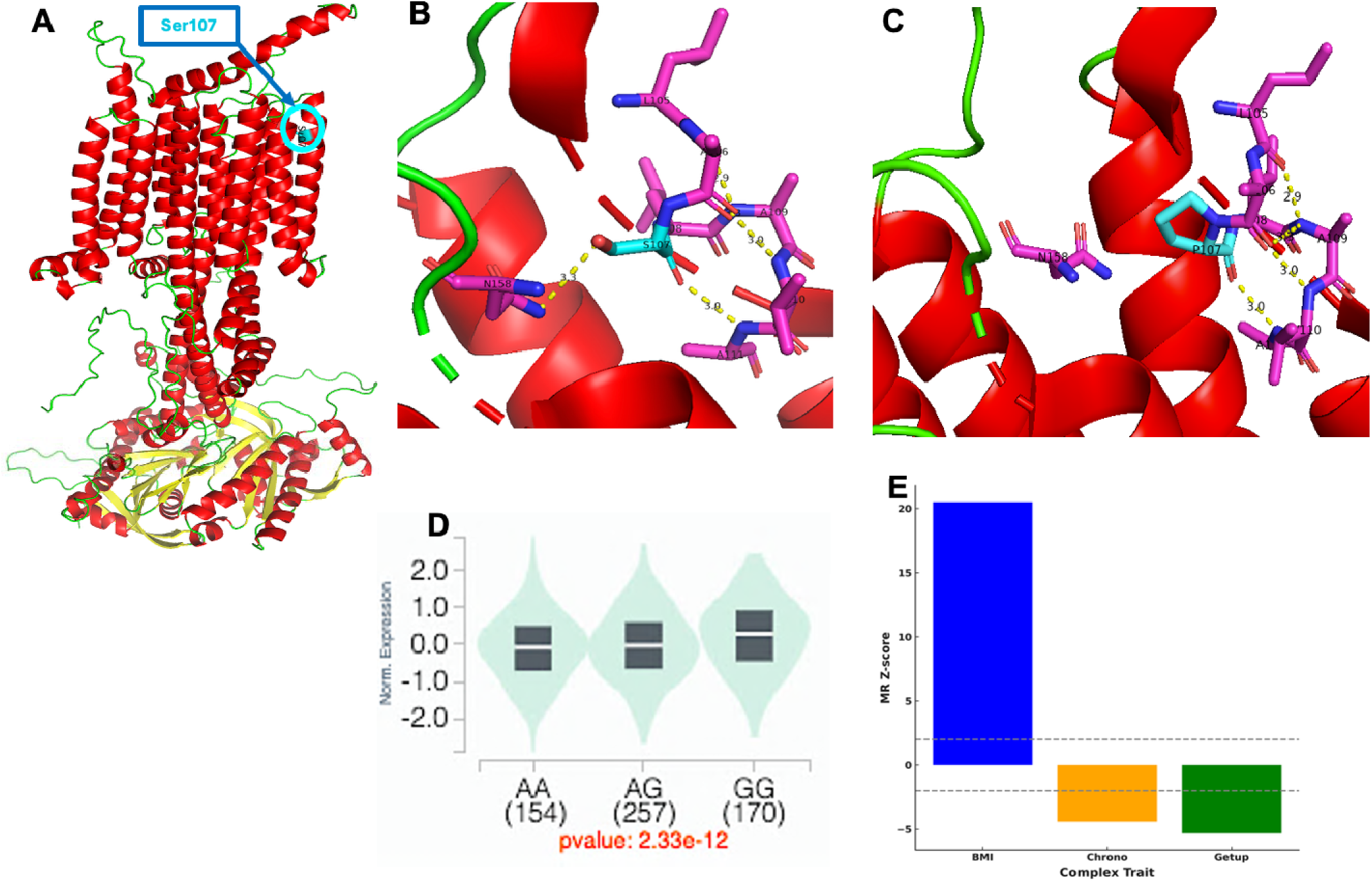
The rs11676272 variant has multi-level functional consequences on protein stability and gene expression. **(A)** The alpha fold predicted the full protein structure of ADCY3, with the location of Serine 107 (S107) highlighted in a transmembrane (TM2) helix. **(B)** A zoomed-in view of Ser107 shows that the hydroxyl oxygen of S107 forms a hydrogen bond (3.3 Å) with the backbone amide nitrogen of Asparagine 158 (N158), and the backbone amide nitrogen of A111 forms a hydrogen bond (3.0 Å) with the hydroxyl oxygen of S107. **(C)** The S107P mutation, which corresponds to the risk G allele, disrupts the stabilizing bond with N158, leading to a predicted increase in protein instability (ΔΔG cartesian = 3.61 kcal/mol). **(D)** In addition to its effect on the protein, the rs11676272 G allele is a significant expression quantitative trait locus (eQTL) associated with higher *ADCY3* expression in subcutaneous adipose tissue (p = 2.33 × 10⁻¹²). **(E)** Mendelian Randomization Wald Ratio method analysis supports a causal relationship between this altered *ADCY3* expression and higher BMI, eveningness chronotype, and more difficulty waking up.

### *Adcy3* is rhythmically expressed in mice adipose tissue, in antiphase to the circadian activator BMAL1

Given *ADCY3’*s link to circadian and adiposity traits in humans, we next investigated the physiological regulation of *Adcy3* in mouse adipose tissue. Using publicly available RNA-seq datasets, we found that *Adcy3* expression was robustly rhythmic in both white (WAT) and brown (BAT) adipose tissue, oscillating in antiphase to the core clock activator *Bmal1* and in phase with the *Per1-3* repressors (**Figures 6A, 6B,** and **Figure S8**). This expression pattern is consistent with the canonical circadian feedback loop, in which BMAL1 and CLOCK activate transcription of the *Per* genes, whose protein products accumulate and inhibit BMAL1/CLOCK activity in a near 24-hour cycle. In contrast, *Adcy3* expression was not rhythmic in the hypothalamus, cerebellum, or liver (**Figure S8**), underscoring the adipose-specific nature of its circadian regulation.

**Figure 6:**
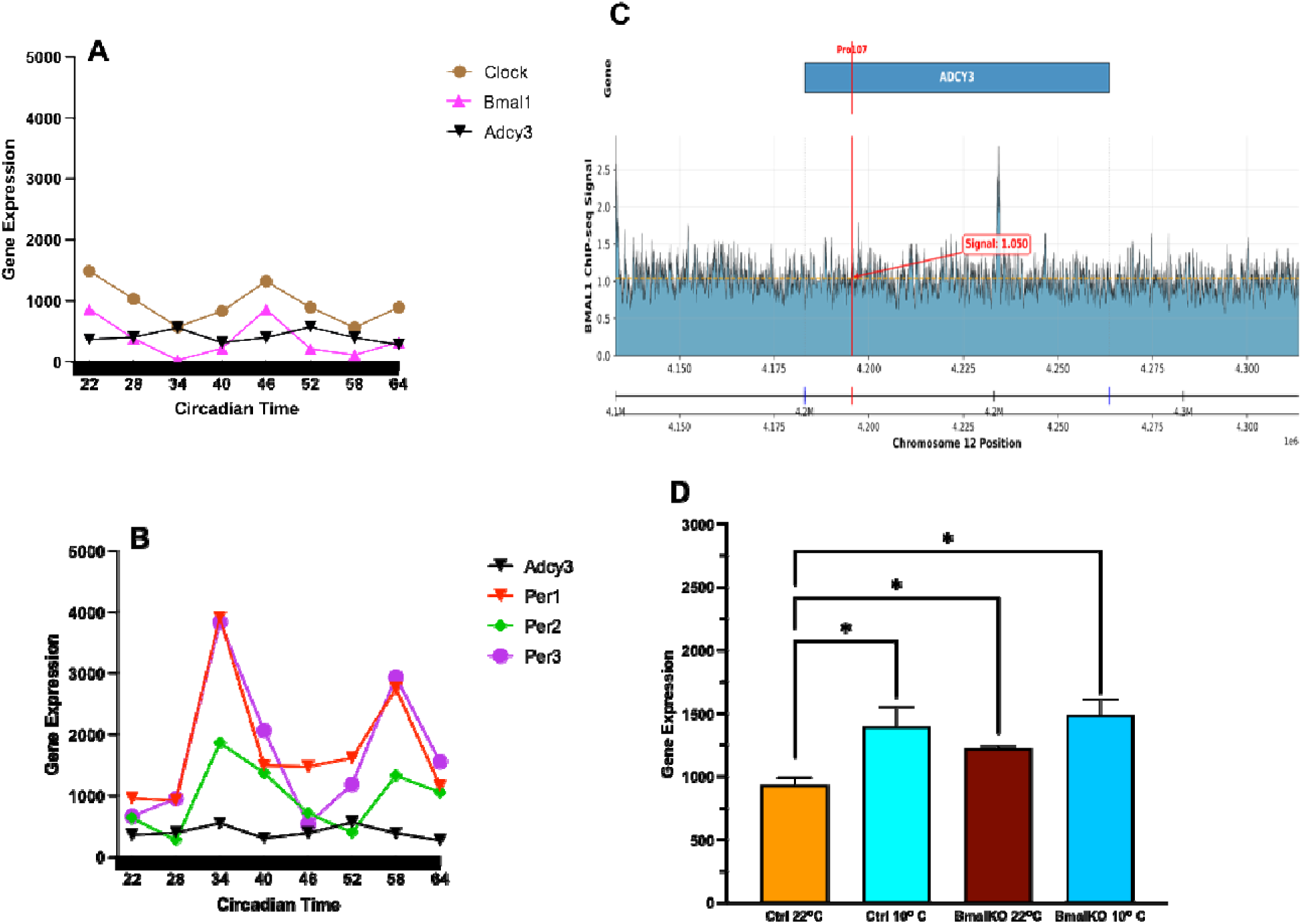
*Adcy3* is a direct, clock-controlled gene regulating thermogenesis in adipose tissue. (**A-B**) *Adcy3* rhythmically oscillates in antiphase to *Clock* and *Bmal1* and in phase with *Per1*/*Per2*/*Per3* over a 42-hour circadian cycle in mouse WAT. (**C**) BMAL1 ChIP-seq in WAT shows binding at the *Adcy3* locus, overlapping the conserved Pro107 site. (**D**) *Adcy3* expression is cold-induced in iWAT of control mice (10°C vs. 22°C), but this response is attenuated by the removal of *Bmal1*. Elevated baseline in knockouts at 22°C suggests derepression in the absence of BMAL1. Data = mean ± SE; *P < 0.05, ANOVA

### BMAL1 binds the conserved Pro107 region of adipose *Adcy3* and regulates its cold-inducible expression

To establish a direct mechanistic link, we analyzed BMAL1 ChIP-seq data from mouse WAT, which revealed binding peaks at the *Adcy3* locus, including one overlapping the conserved orthologous Pro107 site. (**Figure 6C**). These data establish *Adcy3* as a direct transcriptional target of BMAL1. Functionally, we found that the thermogenic induction of *Adcy3* expression by cold exposure was BMAL1-dependent: expression increased significantly in control mice, but was attenuated in *Bmal1* knockout mice, indicating that an intact circadian clock is required for full Adcy3 induction under cold conditions (**Figure 6D**). These findings demonstrate that *Adcy3* is a direct, rhythmic clock output gene in adipose tissue that is functionally integrated with thermogenic pathways.

### *ADCY3* expression in human adipose tissue is a marker of metabolic health

Finally, to assess the clinical relevance of these findings in humans, we analyzed data from the Adipose Tissue Knowledge Portal ^16^. This analysis revealed that *ADCY3* expression is significantly upregulated in human subcutaneous adipose tissue following successful diet-induced weight loss (**Figure 7A**), which is consistent with its proposed role in thermogenic adaptation and energy balance. These two independent observations ^17,18^ provide strong clinical support for our model, suggesting that higher *ADCY3* activity is a hallmark of an improved metabolic state in humans. Together with our genetic and mechanistic data, this positions *ADCY3* as a circadian-regulated metabolic effector that integrates genetic, behavioral, and environmental cues to influence obesity risk.

**Figure 7:**
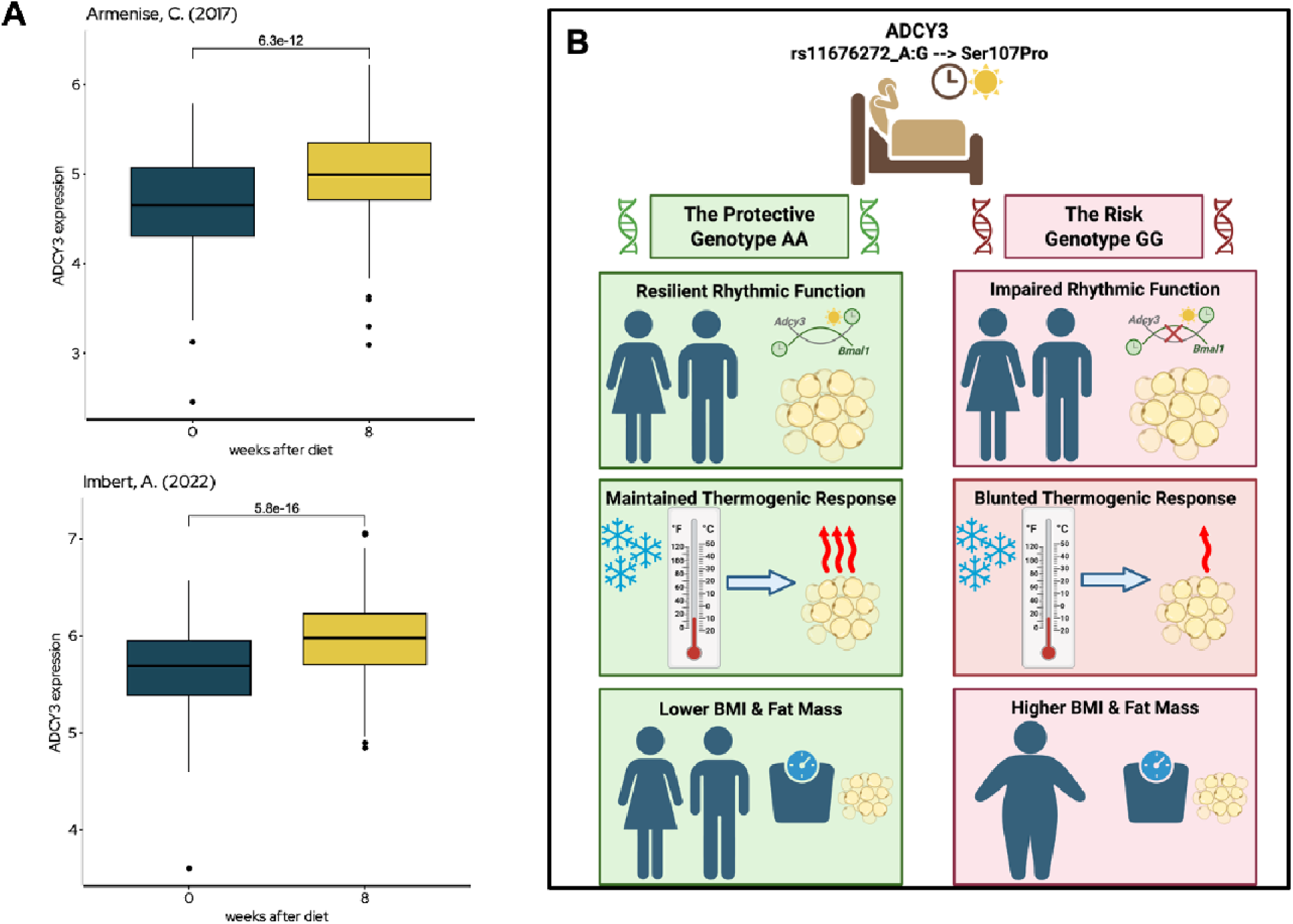
A unified model for *ADCY3* in circadian-metabolic integration and human adipose health. **(A)** *ADCY3* expression increases in subcutaneous adipose tissue following diet-induced weight loss (800-1000kcal/day) in two independent studies (Imbert et al., N = 558; Armenise et al., N = 191), supporting its role in thermogenic recovery and metabolic improvement. (**B**) Schematic model: In AA homozygotes (left), ADCY3 is stable and functional, conferring resilience to circadian strain and supporting rhythmic thermogenesis. In GG homozygotes (right), protein instability and impaired rhythmic response increase vulnerability to lifestyle stressors and promote adiposity.

## DISCUSSION

Obesity arises from a complex interplay of genetic factors, behavioral influences, and tissue-specific physiology. This study identifies the circadian clock-associated gene *ADCY3* as a key genetic mediator of the metabolic consequences of lifestyle factors. We provide evidence from large-scale human genetics and mechanistic mouse studies that a common missense variant in *ADCY3*, rs11676272, interacts with difficulty waking up to significantly increase an individual’s risk of obesity. Our findings reveal a sex-dimorphic effect, a novel mechanism involving protein destabilization with compensatory transcript upregulation and a direct link to thermogenic regulation in adipose tissue, establishing *ADCY3* as a critical node at the interface of circadian biology and metabolic health.

A central finding of our work is the potentiation of genetic risk by a behavioral factor. The obesogenic effect of the G risk allele was most pronounced in individuals who reported difficulty waking up in the morning, a behavioral marker that correlates with BMI. To test the specificity of this finding, we examined other, broader measures of circadian life. Importantly, we observed no significant obesogenic interaction for innate chronotype preference or for the occupational category of night shift work. This crucial distinction suggests that the biological pathway involving *ADCY3* is not uniformly sensitive to all forms of circadian disruption. Rather, our results indicate that its vulnerability is selectively unmasked by increased adiposity, due to difficulty awakening in the morning.

The functional impact of the Ser107Pro mutation underscores the regulatory consequences of rs11676272. Structural modeling shows that the Proline substitution leads to the loss of a stabilizing polar bond and predicts significant destabilization of the ADCY3 protein structure. While this may not impair catalytic function directly, it could influence ADCY3’s role in signaling complexes or membrane dynamics, consistent with prior in-vitro assays showing a modest, though non-significant, reduction in cAMP production in Hek293 cell line ^19^. Notably, the variant also resides within an active promoter and enhancer region in adipose tissue and is associated with a paradoxical increase in *ADCY3* expression. This raises the key possibility that rs11676272 functions as a dual-threat variant, simultaneously altering protein structure while its cis-regulatory effect drives a compensatory increase in transcript levels that is ultimately insufficient to restore normal protein function. This complexity is compounded by tissue-specific isoform usage; we find that adipose tissue expresses a distinct set of ADCY3 transcripts not found in other key metabolic organs, providing a compelling mechanism by which the rs11676272 sQTL can exert its tissue-restricted effects.

Our findings in mice provide a direct physiological context for this mechanism. We show that *Adcy3* is a rhythmic gene specifically in adipose tissues, oscillating in antiphase to its direct transcriptional regulator, BMAL1. Importantly, BMAL1 ChIP-seq reveals a binding peak exactly over the murine Adcy3 segment orthologous to the human Ser107/Pro107 codon, indicating that the same cis-element harboring the missense variant is itself a bona-fide clock-controlled enhancer. This positions *Adcy3* as a direct, rhythmic output of the core clock machinery in fat, tasked with integrating circadian signals with environmental cues. Furthermore, we establish a functional link to energy expenditure, demonstrating that the induction of *Adcy3* by a thermogenic stimulus (cold) is dependent on a functional adipose clock. This regulatory mechanism parallels prior findings at the *FTO* locus, where obesity-associated variants regulate thermogenesis in a tissue-autonomous manner ^20^. Our discovery that full-length *Adcy3* oscillates introduces a crucial circadian dimension to prior ‘rheostat’ models of its function, such as that of the truncated isoform *Adcy3-at* ^21^, suggesting that the temporal gating of *ADCY3* expression is a key mechanism for coordinating thermogenic sensitivity with daily metabolic rhythms.

The role of *ADCY3* as a circadian-regulated metabolic effector involved in thermogenic adaptation is further illuminated by its place in human history, which speaks to the broader debate on the origins of obesity-related genetic variation. This debate often centers on two competing ideas: the ‘thrifty gene’ theory ^22^, which posits that variants were positively selected for energy storage in environments of food scarcity, and the ‘drifty gene’ theory ^23^, which argues that variants accumulated neutrally after predation pressures on fat regulation were relaxed. While neither has been definitively proven, our findings for *ADCY3* align more closely with a model of positive, adaptive selection akin to the thrifty gene concept. Specifically, the protective A allele of rs11676272, which we associate with favorable metabolic outcomes in humans, is more prevalent in non-equatorial populations and exhibits strong signatures of recent positive selection. Its associated gene, *ADCY3*, plays a critical role in thermogenic regulation in mice, further supporting its relevance to cold-adaptive energy balance. This pattern is consistent with prior studies showing that metabolic variants often bear signatures of selection linked to lifestyle and energy balance throughout human history ^24–26^. This suggests that the variant may have conferred an adaptive metabolic advantage in response to specific environmental pressures, such as colder climates. The A allele may have conferred an adaptive metabolic advantage in response to colder climates, consistent with the hypothesis that seasonal or climate-related thermal stress can shape long-term thermogenic programming. Supporting this idea, recent human research by Yoneshiro et al. showed that individuals conceived during colder seasons exhibited higher brown adipose tissue (BAT) activity and greater thermogenic capacity later in life ^27^. Taken together, the population genetics of rs11676272 and its functional role in BMAL1-dependent thermoregulation strongly suggest that *ADCY3* is a gene that has been shaped across human history to fine-tune energy expenditure in response to environmental demands, providing a modern example of adaptive metabolic genetics.

Our findings consistently point to the conclusion that high levels of functional ADCY3 activity are a key feature of a healthy metabolic state in adipose tissue. This principle is supported by two independent lines of evidence. First, our genetic data reveals that the protective A allele of rs11676272, which is predicted to produce a more stable ADCY3 protein, is robustly associated with lower BMI and fat mass. Second, and in perfect alignment with this, independent clinical data show that successful diet-induced weight loss triggers a significant physiological upregulation of *ADCY3* expression in human adipose tissue ^17,18^. This convergence of genetic and clinical evidence solidifies the importance of the peripheral adipose pathway and establishes *ADCY3* as a key marker and potential mediator of metabolic health in humans.

While our data strongly support a peripheral adipose-centric mechanism, we acknowledge the well-established role of *ADCY3* in the central nervous system. *ADCY3* is highly expressed in the brain, and rare, severe loss-of-function mutations were associated with monogenic obesity, highlighting its critical function in central energy homeostasis pathways ^19^. Thus, it is plausible that the pleiotropic effects of this common variant reflect a combined influence in both central and peripheral tissues. In the brain, *ADCY3* localizes to primary cilia of hypothalamic nuclei, including the arcuate nucleus (ARC), suprachiasmatic nucleus (SCN), and paraventricular nucleus (PVN), which are involved in metabolic and circadian regulation ^28–30^. The SCN, recognized as the master circadian clock, coordinates daily rhythms in physiology and behavior ^31–34^. Notably, glutamate stimulation of the SCN increases BAT thermogenesis and core body temperature in rats, demonstrating that central circadian circuits can modulate peripheral energy expenditure ^35^. Furthermore, the melanocortin-4 receptor (MC4R), a Gs-coupled GPCR involved in appetite suppression, co-localizes with ADCY3 at PVN neuron cilia, and disruption of either protein’s ciliary localization impairs energy homeostasis ^29^. Selective inhibition of ADCY3 signaling at MC4R neuronal cilia increases food intake and body weight ^29^, underscoring the importance of ADCY3-mediated cAMP signaling in central metabolic control. Future studies will be necessary to elucidate the relative contributions of central versus adipose ADCY3 signaling to pleiotropic trait associations.

A limitation of our work is that the waking-difficulty item is based on a single self-report question; objective measures such as polysomnography or actigraphy would offer more granular insight into circadian arousal phenotypes. Additionally, the African and South Asian subsets were underpowered for interaction analyses; therefore, future studies with larger multi-ancestry samples will be necessary to evaluate population-specific effects.

Notably, our findings for ADCY3 do not exist in isolation. Other pleiotropic loci identified in our genome-wide pleiotropy screen, such as CELF1, are also known to regulate thermogenic output, albeit through different post-transcriptional mechanisms ^36,37^. This convergence suggests a broader principle in which multiple circadian-metabolic loci modulate energy expenditure through complementary molecular pathways.

In conclusion, we identify ADCY3 as a key integrator of circadian and metabolic signals. We show that a common variant interacts with difficulty awakening to increase obesity risk, linking behavioral circadian strain to adipose thermogenesis through both molecular and physiological mechanisms. These findings establish a new framework for understanding gene–environment interactions in metabolic disease and highlight ADCY3 as a tractable target for rhythm-based interventions aimed at improving metabolic resilience and reducing obesity risk in genetically susceptible individuals.

## RESOURCE AVAILABILITY

### Lead Contact

Further information and requests for resources should be directed to and will be fulfilled by the Lead Contact, Richa Saxena

### Materials Availability

This study did not generate new, unique materials.

### Data and Code Availability

All publicly available GWAS summary statistics used in this study can be accessed via the UK Biobank and GIANT Consortium websites and the Knowledge Portal at https://hugeamp.org/. GTEx v8 expression data is available at https://gtexportal.org/.

## ACKNOWLEDGEMENTS

This research has been conducted using the UK Biobank Resource (UK Biobank application number 6818). We would like to thank the participants and researchers from the UK Biobank study and Genetic Investigation of ANthropometric Traits (GIANT) Consortium. C.T. is supported by BWF G-1022367. J.M. is supported by a Humboldt Professorship of the Alexander von Humboldt Foundation. J.M. acknowledges funding by the Deutsche Forschungsgemeinschaft (DFG) through SFB1423 (421152132), SFB 1664 (514901783), TRR 386 (514664767), and SPP 2363 (460865652). J.M. is supported by the Federal Ministry of Education and Research (BMBF) through the Center for Scalable Data Analytics and Artificial Intelligence (ScaDS.AI), through the German Network for Bioinformatics Infrastructure (de.NBI), and through the German Academic Exchange Service (DAAD) via the School of Embedded Composite AI (SECAI 15766814). Work in the Meiler laboratory is further supported through the National Institute of Health (NIH) through R01 HL122010, R01 DA046138, R01 AG068623, R01 LM013434, S10 OD016216, S10 OD020154, S10 OD032234; R.S. is supported by NIH R01 DK107859, NIH R01 DK102696, NIH R01 HL146751 and MGH Research Scholar Fund. J.L. is supported by NIH/NHLBI K01 HL136884 and NIH/NHGRI R01 HG012810.

## AUTHOR CONTRIBUTIONS

C.T. and R.S. designed the study; J.L. conducted chronotype GWAS for the UKB cohort; C.T. acquired the BMI GWAS summary statistics. C.M. and J.M. built the protein structural biology pipeline, and C.T. conducted the remaining human genetics and in-vivo periodicity analyses. All co-authors participated in acquiring, analyzing, and interpreting the data. C.T. wrote the manuscript, and all co-authors reviewed and edited the manuscript before approving its submission. R.S. is the guarantor of the work and, as such, has full access to all the data in the study and takes responsibility for the integrity of the data and the accuracy of the data analysis.

## DECLARATION OF INTEREST

The authors have declared no competing interests.

## SUPPLEMENTAL INFORMATION

Supplemental Figures. Figure S1-S9 Supplemental Tables: Table S1-S9.

## STAR METHODS

Detailed methods are provided in the online version of this paper and include the following:

## KEY RESOURCES TABLE RESOURCE AVAILABILITY

- Lead contact
- Materials availability
- Data and code availability

## EXPERIMENTAL MODEL AND SUBJECT DETAILS

- Human genetic datasets
- Animal models and publicly available RNA-seq datasets METHOD DETAILS
- Genome-wide pleiotropy and colocalization analysis of circadian preference and BMI
- Functional Annotation and Causal Inference
- Tissue-specific colocalization and causal inference
- Phenome-wide Association Study
- UK Biobank Genetic and Phenotypic Model Analysis
- Protein structural modeling
- ADCY3 circadian rhythmicity in animal models
- Human ADCY3 Tissue-Specific Isoform Usage QUANTIFICATION AND STATISTICAL ANALYSIS
- Genome-Wide Pleiotropy Analysis
- Colocalization Analysis
- Linear mixed-model analyses (UK Biobank)
- Mendelian Randomization
- Protein Structural Modeling and Stability Analysis
- Circadian Expression Analysis
- Tissue-Specific Isoform Regulation and BMAL1 Binding Motif Analysis
- Statistical Software and Significance Thresholds

## KEY RESOURCES TABLE

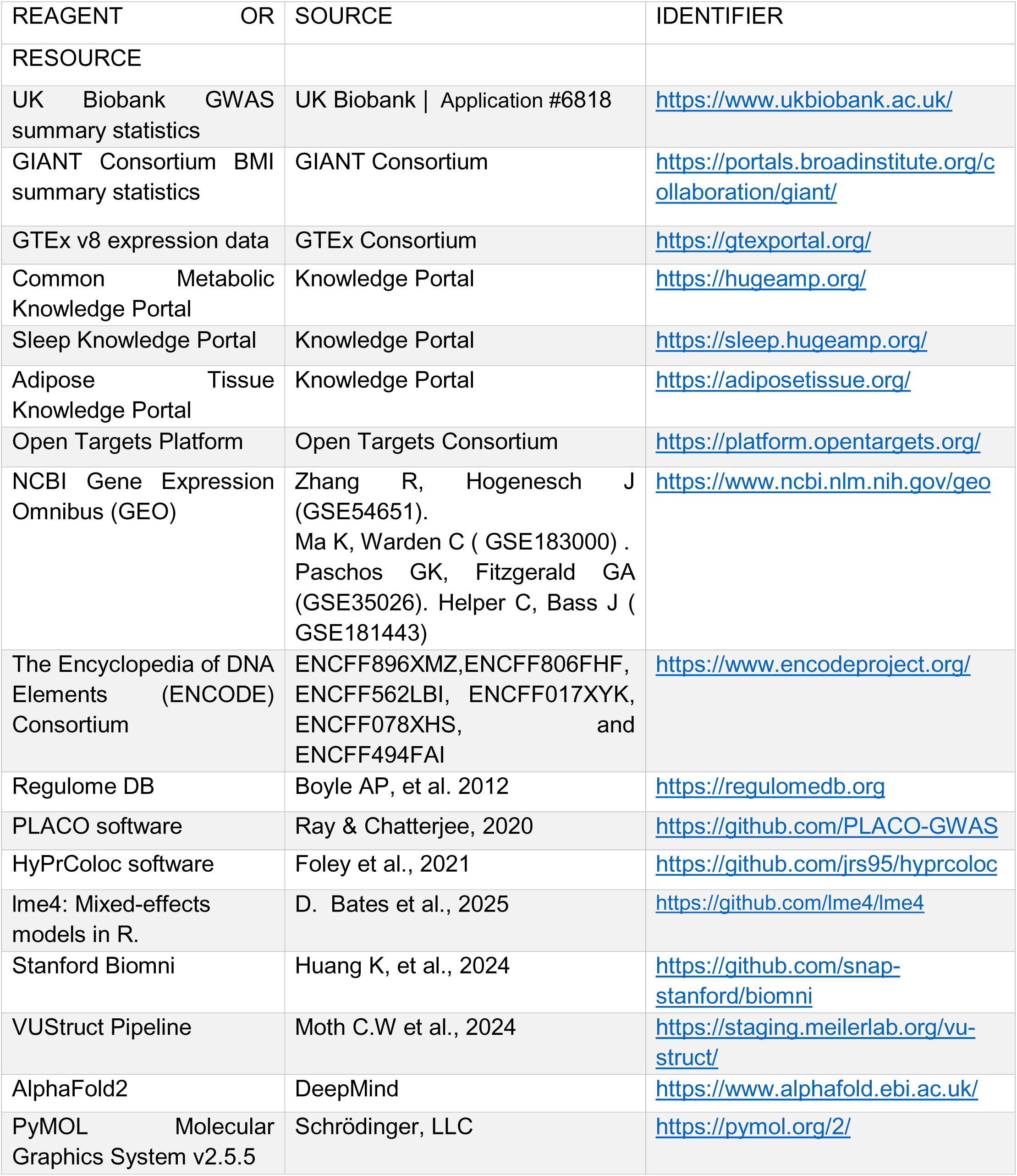

## RESOURCE AVAILABILITY

### Materials Availability

This study did not generate new, unique materials.

### Data and Code Availability

All publicly available GWAS summary statistics used in this study can be accessed via the UK Biobank and GIANT Consortium websites and the Knowledge Portals at https://hugeamp.org/ | https://sleep.hugeamp.org/ | https://adiposetissue.org/ . GTEx v8 expression quantitative loci data is available at https://gtexportal.org/. The publicly available mouse RNAseq and ChIP-Seq data are available at https://www.ncbi.nlm.nih.gov/geo . The Human Histone ChIP-Seq data are available https://www.encodeproject.org/. The Transcription Factors ChIP-Seq data are available in https://regulomedb.org

## EXPERIMENTAL MODEL AND SUBJECT DETAILS

### Human Samples

We analyzed GWAS summary statistics from the UK Biobank (UKB), which includes ∼500,000 individuals. Chronotype summary statistics were derived from self-reported morningness phenotype (N = 449,734) ^4^, and BMI data were drawn from a meta-analysis combining UKB and the GIANT consortium (total N = 694,649) ^14^. A GWAS for ease of waking up in the morning was conducted de novo in UKB with adjustments for age, sex, genotyping array, and 10 genetic principal components (N = 451,872) under the UKB application # 6818. No new human subjects were recruited; all analyses used de-identified, publicly available summary-level data

### Animal Models

Circadian expression and regulatory data were obtained from C57Bl6 male mice using publicly available datasets. Gene expression analyses utilized the following RNA-seq datasets: GSE54651, for tissues harvested under constant darkness following circadian entrainment; GSE183000, for data from mice exposed to cold (10°C) or ambient temperature (22°C), with wild-type and Bmal1 knockout models. To assess direct transcriptional regulation by the core clock, we analyzed a BMAL1 ChIP-seq dataset from mouse inguinal white adipose tissue (iWAT) (GSE181443). All experiments and datasets were previously published.

## METHOD DETAILS

### Genome-wide pleiotropy and colocalization analysis of circadian preference and BMI

To identify shared genetic signals between circadian preference and BMI, we used PLACO (Pleiotropic Analysis under Composite Null Hypothesis v0.1.1), a statistical approach that detects pleiotropic loci based on summary-level GWAS data from two traits ^12,38^. We harmonized effect alleles, removed SNPs with mismatched alleles, and filtered datasets for minor allele frequency (MAF) > 1%. The Pearson correlation of Chrono-BMI and Getup-BMI Z-scores were 0.0758 and 0.0871, respectively. A total of 67,449 SNPs were available for Chrono-BMI pleiotropy analysis and 41,808 SNPs for Getup-BMI pleiotropy. Given the large effect sizes in BMI GWAS, applying the recommended Z² > 80 filtering would have disproportionately removed variants with strong pleiotropic effects. Since the GWAS summary statistics have overlapping samples/individuals from the UK-Biobank, we decorrelate the Z-scores using the correlation matrix. To ensure the retention of biologically meaningful SNPs, we validated our PLACO findings through HyPrColoc (Hypothesis Prioritization in Multi-Trait Colocalization v1.0) ^13^ , a robust method prioritizing shared causal variants across multiple traits. For HyPrColoc, we used the default parameters of prior.2 = 0.98, bb.alg = F. To visualize pleiotropic loci, we generated Manhattan and Q-Q plots using the qqman R package ^39^ All the regional association plots were generated using LocusZoom ^40^ from the GWAS summary statistics for circadian preference and BMI.

### Functional Annotation and Causal Inference

We annotated significant variants using FUMA (v1.5.2) ^41^. Lead SNPs were identified using a P-value threshold of 5 × 10⁻ , and linkage disequilibrium (LD) was assessed using the UKB British 10K reference panel (LD R² threshold: lead SNPs = 0.6, secondary SNPs = 0.1, positional mapping window = 10 kb, locus merge distance = 250 kb).

To investigate regulatory functions, we integrated GTEx v8 cis-eQTLs ^42^ from VannoPortal ^43^ and Roadmap Epigenomics data ^44^ to identify tissue-specific regulatory effects. The human subcutaneous adipose tissue ChIP-Seq for enhancer epigenetic markers H3K4me1 ( ENCFF896XMZ, ENCFF806FHF, and ENCFF562LBI) and promoter marker H3K4me3 ( ENCFF017XYK, ENCFF078XHS, and ENCFF494FAI) were obtained from ENCODE, and transcription factor binding data from RegulomeDB45.

### Tissue-specific colocalization and causal inference

To assess whether pleiotropic signals reflect shared genetic regulation, we applied HyPrColoc to evaluate colocalization between PLACO-identified loci and tissue-specific eQTLs. We prioritized loci within ADCY3, CELF1, FTO, and TFAP2B based on their pleiotropic effects on circadian traits and BMI. HyPrColoc estimates posterior probability (PP) for colocalization, identifying candidate causal SNPs contributing to the total colocalization signal.

To assess the causality and directionality of ADCY3 expression on complex traits, we performed Mendelian Randomization (MR) analysis focused on the missense variant rs11676272 using the TwoSampleMR R package (v0.6.14) ^46,47^. The eQTL effect of rs11676272 on ADCY3 expression in subcutaneous adipose tissue (SAT) was used as the exposure, and GWAS summary statistics for BMI, morningness chronotype (Chrono), and ease of waking up (Getup) were used as outcomes. Causal estimates were derived using the Wald ratio method, which is optimal when a single variant is used as an instrumental variable. The MR Z-score was computed as MR Beta / MR SE. **Phenome-wide Association Study**

To explore additional associations, we used OpenTargets ^48^ to conduct a phenome-wide association study (PheWAS) using GWAS summary statistics from FinnGen, UK Biobank, and the GWAS Catalog.

### UK Biobank Genetic and Phenotypic Model Analysis

To dissect the effects of the lead variant rs11676272, we performed detailed analyses in the UK Biobank cohort. All linear regression models testing the main effect of genotype, sex-dimorphism, and gene-by-behavior interactions were adjusted for age, sex, and the first 10 genetic principal components to control for population stratification. Analyses were performed in R (v4.1.0). Full model summaries are available in Table S8.

### Protein structural modeling

We performed calculations and visualization on the ADCY3 full-length AlphaFold 2 ^49^ model of UniProt ID O60266-1, as well as 37% identity Swiss homology Models templated on CryoEM structures 8sl4.pdb (ADCY5 positions 51 to 1124) and 8buz.pdb (ADCY8 positions 59 to 1114 positions). We used the ΔΔG Cartesian ^50,51^ protocol as implemented in the VUStruct pipeline ^52^ to predict whether S107P might negatively impact ADCY3’s free energy of folding. The PyMOL Molecular Graphics System, Version 2.5.5, Schrödinger, LLC, was used to visualize ADCY3 protein structure with a focus on changes to the hydrogen bonding network around Serine 107 on the mutation to Proline.

### ADCY3 circadian rhythmicity in animal models

To assess endogenous circadian rhythmicity, we analyzed publicly available RNA-seq data from C57Bl/6 male mice (GSE54651) ^53^, where 6-week-old mice were entrained to a 12h:12h light:dark schedule for 1 week, then placed in constant darkness, and tissues were harvested at circadian time CT 22-64 every 6 hours ^53^. RNA was extracted using Qiagen RNeasy kit, and Illumina HiSeq 2000 was used for sequencing. DeSeq2 was used for normalization. We used the MetaCycle R package ^54^ to assess the rhythmicity of gene expression from white adipose, brown adipose, hypothalamus, cerebellum, liver, and aorta. For periodicity analysis in MetaCycle, we used Meta2d method that integrates the ARS (Autoregressive Spectral analysis), the JTK_CYCLE (Jonckheere-Terpstra-Kendall Cycle), and the LS (Lomb-Scargle periodogram) to analyze rhythms in time-series data^54^. This integrated approach improves detection power by combining complementary algorithms for rhythmicity in time-series data without replicates. GraphPad Prism (v.10.2.3) was used to generate the rhythmic gene expression figures. Due to ADCY3 adipose-specific rhythmicity, we used openly available RNAseq data from NCBI Geo GSE183000 for C57Bl/6 mice inguinal white adipose tissue (iWAT) ^55^. The mice were kept in a 12:12 light/dark cycle. For adipose-specific Bmal1 target, Bmal1fl/fl and Prx1-Cre transgenic mice alongside CRISPR-engineered ROSA-26 Bmal1 knock-in and knockout mice. Then, the 10-12 week-old male mice were placed in an ambient temperature of 22°C and a cold temperature of 10°C with a 2-week acclimation period, and the iWATs were collected at ZT3. Total RNA was extracted using Trizol, and Illumina HiSeq 2500 was used for sequencing. GraphPad Prism was used for the analysis and to generate the figures. Additionally, the binding of BMAL1 to ADCY3 was further confirmed in mice using iWAT openly accessible ChipSeq data (GSE181443) ^56^, and we used Biomni Stanford ^57^ to visualize the BMAL1 peak; signal intensities were calculated as mean values across 500-bp genomic bins spanning the ADCY3 locus (chr12:4,133,103-4,313,525, GRCm39 assembly) with 50-kb flanking regions . We then query the *ADCY3 homo sapiens* sequence from the NCBI (ADCY3/NM_001377130.1/NP_001364059.1) to scan for the canonical *Bmal1*-specific E-box motif CACGTG ^58^, and we identified 3 E-box motifs in human.

### Human ADCY3 Tissue-Specific Isoform Usage

We used GTex v.8 transcript browser to assess tissue-specific isoform expression of ADCY3 across different tissues, as the splicing regulation and isoform length might explain tissue-specific rhythmicity, as it might result in the removal of BMAL1 binding motifs.

## QUANTIFICATION AND STATISTICAL ANALYSIS

### Genome-Wide Pleiotropy Analysis

Genome-wide pleiotropy analysis was conducted using PLACO (Pleiotropic Analysis under Composite Null Hypothesis v0.1.1), a statistical framework that identifies shared genetic effects across two traits. PLACO was applied to identify SNPs influencing morning chronotype, ease of waking up, and BMI, with statistical significance determined using a genome-wide significance threshold of P < 5 × 10⁻ . SNP harmonization was performed by aligning effect alleles across summary statistics, and variants with minor allele frequency (MAF) < 1% were excluded. Pearson correlation of Z-scores was calculated to assess the degree of pleiotropy between Chrono-BMI and Getup-BMI, yielding correlation coefficients of 0.0758 and 0.0871, respectively. To validate our pleiotropy findings, we performed a secondary pleiotropy analysis using HyPrColoc, a Bayesian method that prioritizes shared causal variants across multiple traits. HyPrColoc colocalization results were considered significant when the posterior probability (PP) exceeded 0.60, indicating strong evidence of a shared causal genetic signal.

### Colocalization Analysis

Colocalization analysis was performed to determine whether pleiotropic loci exhibit a shared causal effect on gene expression and complex traits. We used HyPrColoc (Hypothesis Prioritization in Multi-Trait Colocalization v1.0) to estimate posterior probabilities (PP) of colocalization between PLACO-identified SNPs and expression quantitative trait loci (eQTLs) in functionally relevant tissues obtained from GTEx v8. Colocalization was performed using a ± 1kb window around each gene of interest based on the GRCh37/hg19 genome build. The genomic regions analyzed were: *ADCY3* (chr2:25,041,038–25,144,106), *CELF1* (chr11:47,486,489–47,588,091), *FTO* (chr16:53,736,875–54,156,853), and *TFAP2B* (chr6:50,785,584–50,816,332). A colocalization signal was considered noteworthy when PP > 0.60, suggesting that the same genetic variant regulates gene expression and the associated trait.

### Linear mixed-model analyses (UK Biobank)

All genotype–phenotype associations were tested with linear mixed models in R (lme4 v1.1-34; lmerTest v3.1-3). For each quantitative trait (BMI, fat mass, % body fat, chronotype score, and morning difficulty waking score), we fitted Trait ∼ Genotype + Age + Sex + Array + PC1–PC10 + (1 | Kinship_ID) where Genotype were coded as a three-level factor (reference = GG; comparison levels AG and AA). Kinship_ID is a random intercept that groups all individuals belonging to the same UK Biobank kinship component, thereby accounting for cryptic relatedness. Sex-stratified models omitted the Sex term; interaction models added the relevant cross-product (e.g., Genotype × Sex or Genotype × GET_UP). Complete model outputs are provided in Table S8

### Mendelian Randomization

Mendelian randomization (MR) was performed to determine whether ADCY3 expression causally influences morning chronotype and BMI. The TwoSampleMR R package (v0.6.14) ^46,47^ was used to integrate eQTL and GWAS data. Given that rs11676272 was the only available genetic instrument for ADCY3 expression in adipose tissue, we applied the Wald ratio method, which estimates causal effects by dividing the variant’s effect size on the outcome by its effect on the exposure.

The eQTL effect of rs11676272 on ADCY3 expression in subcutaneous adipose tissue (SAT) was used as the exposure, and GWAS summary statistics for BMI, morningness chronotype (Chrono), and ease of waking up (Getup) were used as outcomes. Causal estimates were derived using the Wald ratio method, which is optimal when a single variant is used as an instrumental variable. The MR Z-score was computed as MR Beta / MR SE and significance was assessed at P < 0.05.

### Protein Structural Modeling and Stability Analysis

Protein structural analysis was conducted to evaluate the impact of rs11676272 (S107P) on ADCY3 stability. Structural modeling was performed using AlphaFold2 and Swiss homology models, comparing ADCY3 against CryoEM templates of ADCY5 and ADCY8. Structural stability was assessed using Rosetta ddG Cartesian calculations, where ΔΔG values greater than 3 kcal/mol were considered indicative of significant destabilization. The PyMOL Molecular Graphics System (v2.5.5) was used for molecular visualization, specifically examining hydrogen bonding networks, polar interactions, and structural dynamics following the S107P substitution.

### Circadian Expression Analysis

Circadian expression analysis was conducted to assess the rhythmic regulation of ADCY3 in adipose tissue. RNA-seq datasets from mice (C57Bl6, GSE54651, GSE183000) were analyzed using MetaCycle, a statistical tool that integrates ARS (Autoregressive Spectral Analysis), JTK_CYCLE (Jonckheere-Terpstra-Kendall Cycle), and LS (Lomb-Scargle Periodogram) methods to detect oscillatory gene expression patterns. BMAL1-dependent transcriptional regulation was further evaluated using ChIP-Seq data, which revealed BMAL1 binding peaks at the ADCY3 promoter in mouse inguinal white adipose tissue (iWAT).

### Tissue-Specific Isoform Regulation and BMAL1 Binding Motif Analysis

Tissue-specific isoform expression of ADCY3 was analyzed using GTEx v8 isoform data. Differential isoform expression was assessed across multiple tissues, with adipose-specific transcripts absent in the liver, whole blood, hypothalamus, pancreas, and heart. To determine whether alternative splicing affects BMAL1 binding and ADCY3 rhythmicity, we examined the presence of BMAL1-specific E-box motifs (CACGTG) within the ADCY3 promoter region. Three BMAL1-specific E-box binding sites were identified suggesting that BMAL1 directly regulates ADCY3 in a tissue-specific manner.

### Statistical Software and Significance Thresholds

All statistical analyses were performed using R (v4.1.0), Python (v3.9), ggplot2 (v3.5.0), and GraphPad Prism (v10.2.3). The significance threshold for all tests was set at P < 0.05 unless otherwise stated. Full statistical details, including sample sizes, effect sizes, confidence intervals, and P-values, are provided in the figure legends and supplementary tables.

