## Supplementary material for "*ADCY3* Ser107Pro links difficulty awakening in the morning to adiposity through circadian regulation of adipose thermogenesis": Figure S

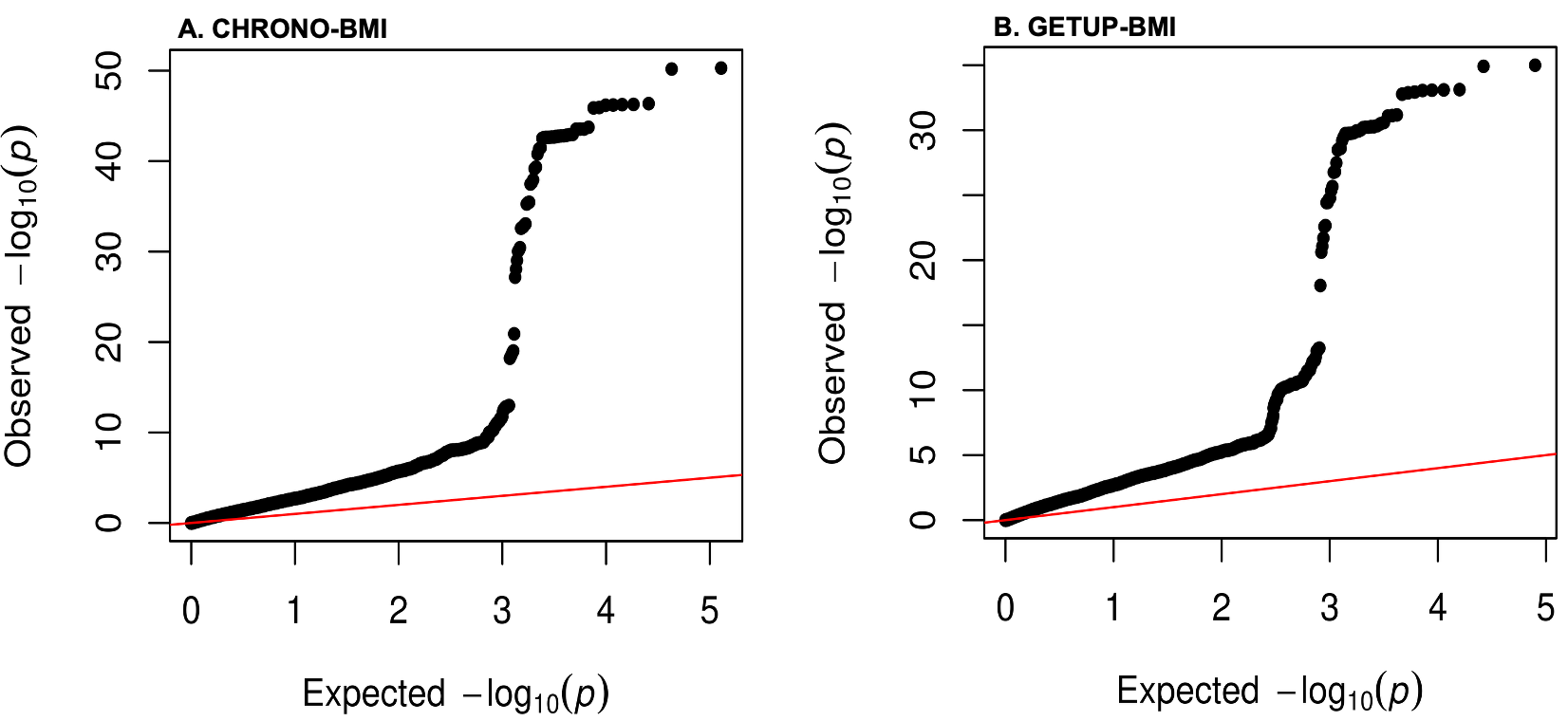

**Figure S1: Quality Control of Genome-wide Pleiotropy Analyses.** Quantile-Quantile (QQ) plots showing observed versus expected –log₁₀(p) values for pleiotropy analyses between (**A**) morningness chronotype and BMI and (**B**) ease of getting up and BMI.

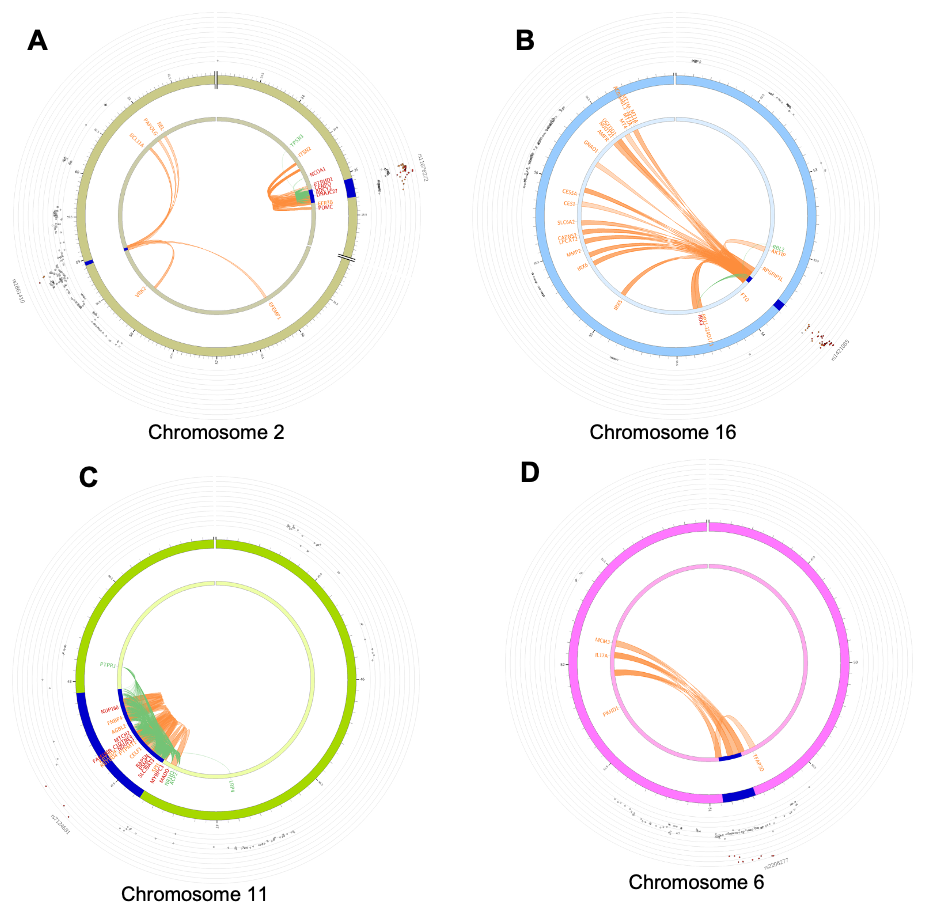

**Figure S2: Regulatory landscape of pleiotropic loci linking circadian and metabolic traits.** Circos plots visualize the functional genomic context of the four top pleiotropic loci identified by genome-wide analysis: (**A**) *ADCY3*, (**B**) *FTO*, (**C**) *CELF1*, and (**D**) *TFAP2B*. For each locus, the outermost ring shows local association p-values from the pleiotropy analysis. Orange arcs indicate chromatin interactions (Hi-C) derived from adipose-derived mesenchymal stem cells. Green arcs represent significant expression quantitative trait loci (eQTLs) in key metabolic and circadian tissues, as summarized in Table S5.

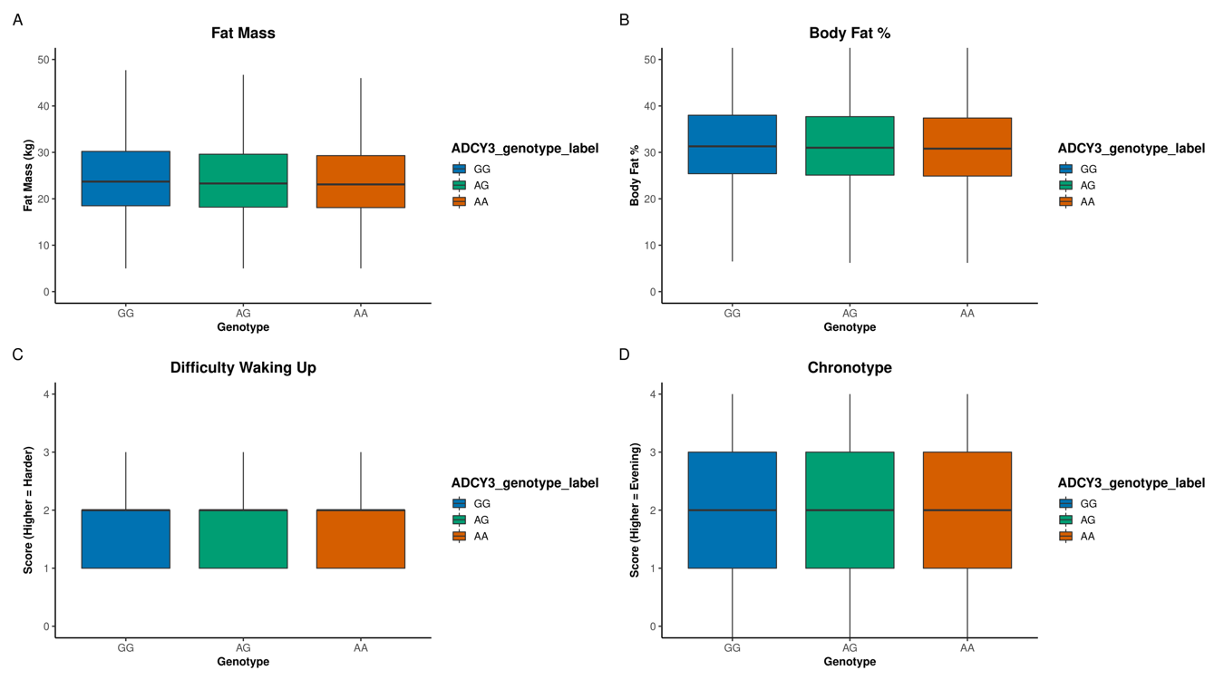

**Figure S3: Main effect of ADCY3 rs11676272 genotype on adiposity and circadian-behavioral phenotypes.** Boxplots show associations between ADCY3 genotype (rs11676272) and (**A**) fat mass, (**B**) body fat percentage, (**C**) difficulty waking, and (**D**) chronotype in UK Biobank Europeans (N= 451,324). The G allele is associated with significantly higher adiposity (e.g., AA vs GG:β = –0.47 kg for fat mass, β = –0.40% for body fat; both P < 2 × 10⁻¹⁶), and more difficulty waking up (AA vs GG: β = –0.016, P = 3.6 × 10⁻⁷). The association with chronotype (AA vs GG: β = –0.016, P = 0.0001) trends toward eveningness but is a smaller magnitude. All models were adjusted for age, sex, PC1–10, and kindship. Boxes show the interquartile range (IQR), center lines denote medians, and whiskers span 1.5 × IQR. Colors: GG (blue), AG (green), AA (orange).

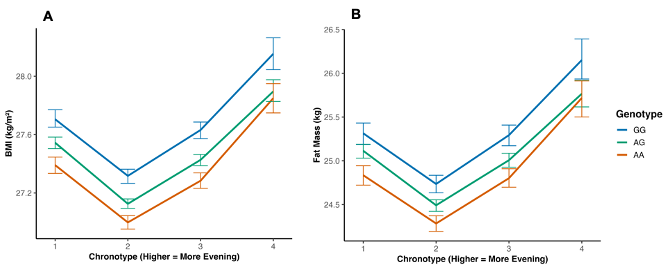

**Figure S4: No significant interaction between *ADCY3* genotype and chronotype preference on adiposity.** Interaction plots show the relationship between ADCY3 rs11676272 genotype and chronotype preference on (**A**) BMI and (**B**) fat mass in UK Biobank Europeans (N ≈ 403,000). Points represent mean values per genotype × chronotype group; error bars denote 95% confidence intervals. Genotype–chronotype interaction terms in linear models were not statistically significant for BMI (AG × Chrono: β = –0.032, P = 0.11; AA × Chrono: β = –0.012, P = 0.60) or fat mass (AG × Chrono: β = –0.067, P = 0.087; AA × Chrono: β = –0.014, P = 0.76). All models adjusted for age, sex, PC1–10, and kindship. Colors: GG (blue), AG (green), AA (orange).

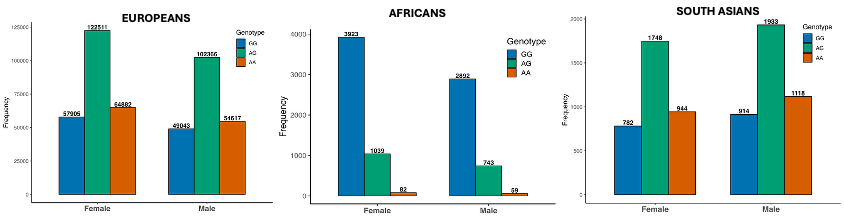

**Figure S5: Sex- and ancestry-specific carrier counts for ADCY3 rs11676272 in the UK Biobank.** Bar plots show the number of individuals with each rs11676272 genotype (GG, AG, AA) stratified by sex in (**A**) Europeans (EUR), (**B**) Africans (AFR), and (**C**) South Asians (SAS). Participant carrier counts are displayed above each bar. The plots highlight the large disparity in sample sizes across ancestries and genotype groups, particularly the limited number of AA homozygotes in non-European populations, which constrains power for interaction and stratified analyses in these groups.

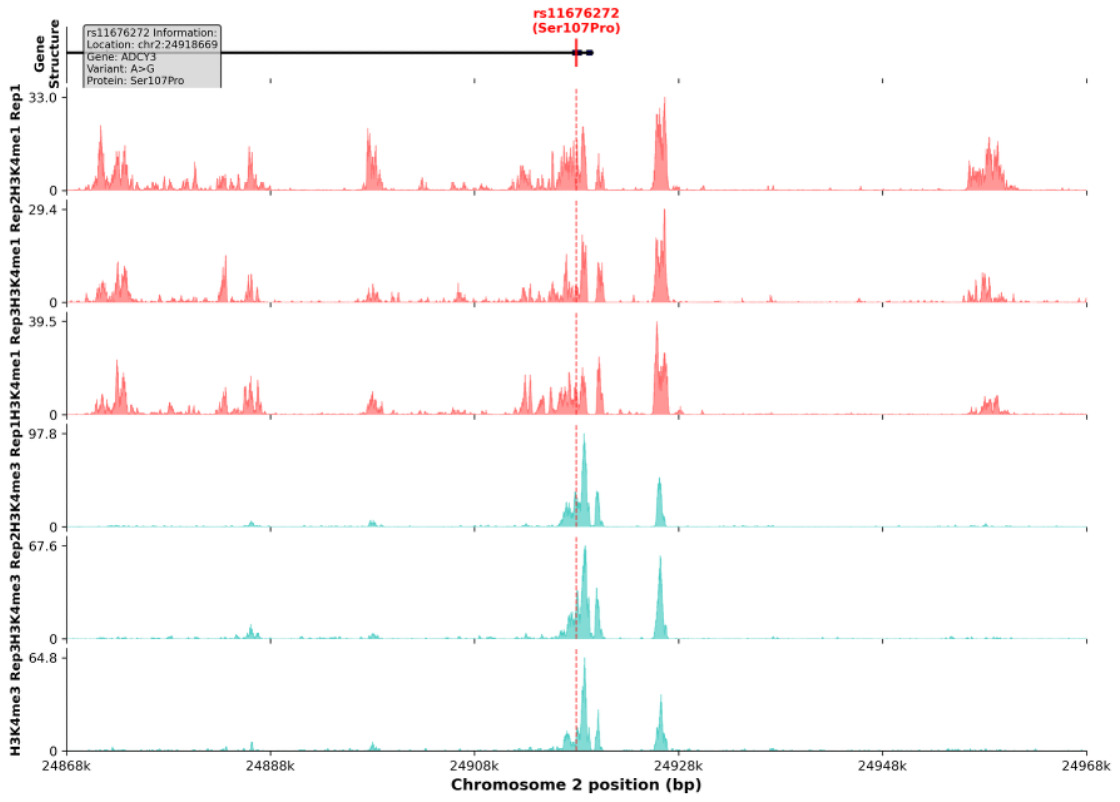

**Figure S6: The rs11676272 region overlaps active regulatory chromatin in human subcutaneous adipose tissue.**

Genome browser visualization showing H3K4me1 and H3K4me3 ChIP-seq signals at the rs11676272 locus in the ADCY3 gene. The analysis spans chr2:24,868,669-24,968,669 (100kb window) and includes: (**Top**) Gene structure track showing ADCY3 exons (dark blue rectangles) with rs11676272 position marked (red line); (**Tracks 2-4**) H3K4me1 ChIP-seq signals from three biological replicates indicating enhancer activity; (**Tracks 5-7**) H3K4me3 ChIP-seq signals from three biological replicates indicating promoter activity. The variant lies within a region of strong chromatin activity for both markers, suggesting it may act as a cis-regulatory element in adipose tissue..

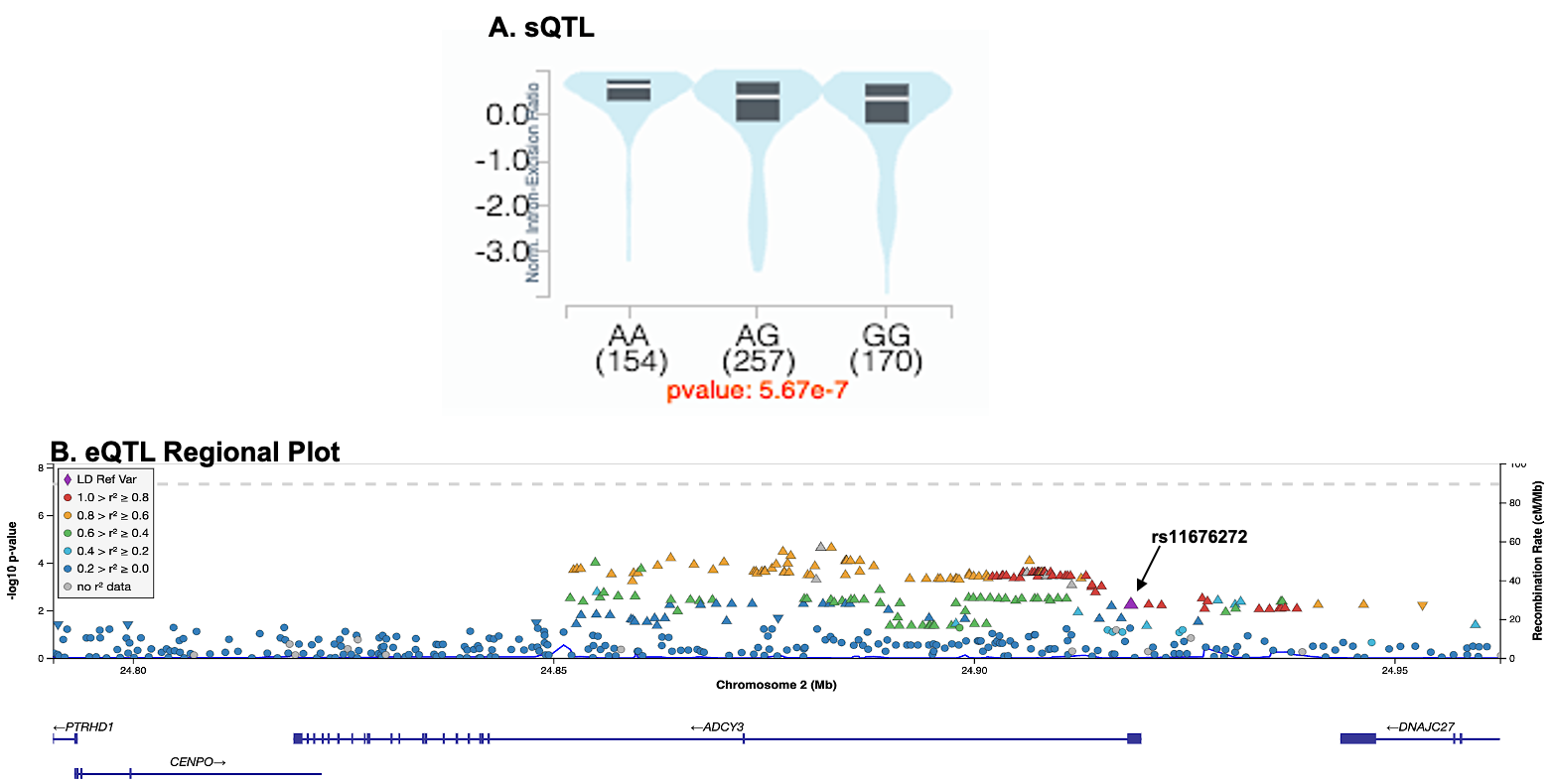

**Figure S7: rs11676272 is a splicing and expression QTL for ADCY3 in human adipose tissue.** (**A**) Violin plot of splicing QTL (sQTL) data from GTEx v8 showing the effect of rs11676272 genotype on intron inclusion in ADCY3 transcripts in subcutaneous adipose tissue (p = 5.67 × 10⁻⁷). The G allele (risk) is associated with reduced intron retention, consistent with altered isoform composition. (**B**) Regional association plot of cis-eQTLs in subcutaneous adipose tissue confirms rs11676272 as the lead SNP regulating ADCY3 expression. The y-axis indicates –log₁₀(p-value) for expression association; SNPs are color-coded by LD (r²) with rs11676272 using the 1000 Genomes EUR reference panel.

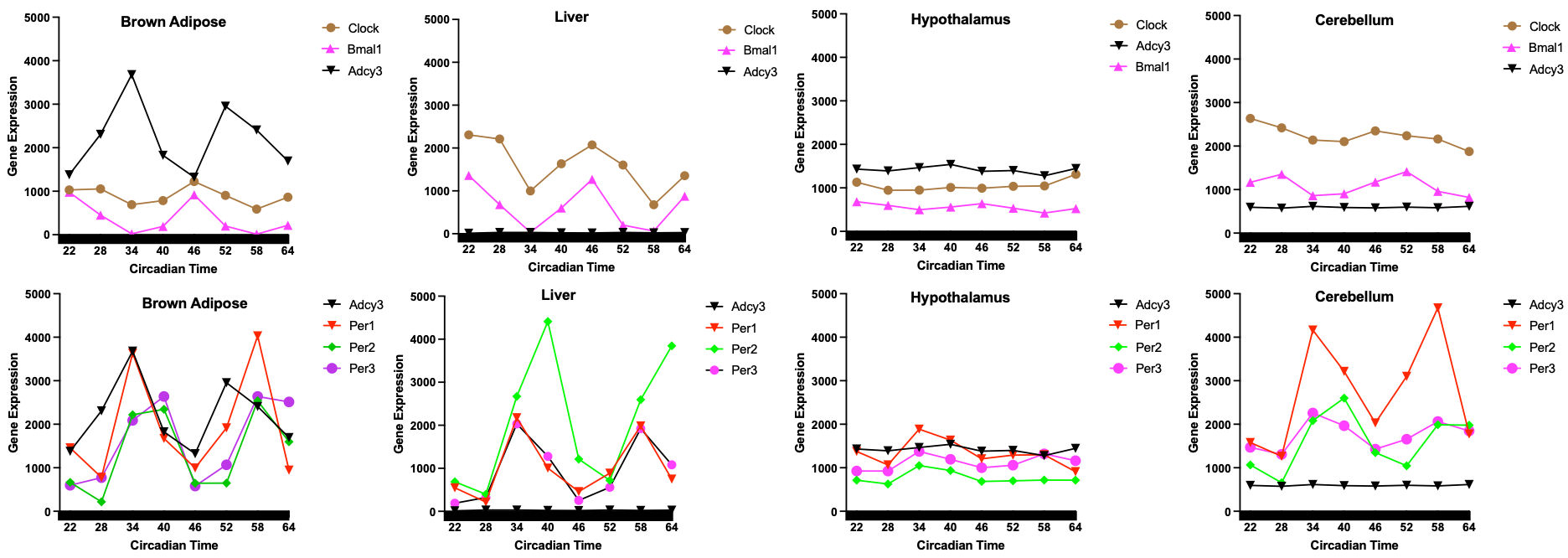

**Figure S8: *Adcy3* exhibits tissue-specific circadian rhythmicity in mouse adipose tissues.**
Time-series RNA-seq expression profiles of *Adcy3* and core clock genes (*Bmal1, Per1, Per2, Per3*) across a 42-hour circadian cycle in mouse tissues, based on MetaCycle analysis. *Adcy3* expression displays significant rhythmicity in white (WAT) and brown adipose tissue (BAT), oscillating in antiphase to *Bmal1* and in phase with *Per1-3* genes. No significant rhythmicity was observed in the liver, hypothalamus, or cerebellum. These results underscore the adipose-specific circadian regulation of *Adcy3*.

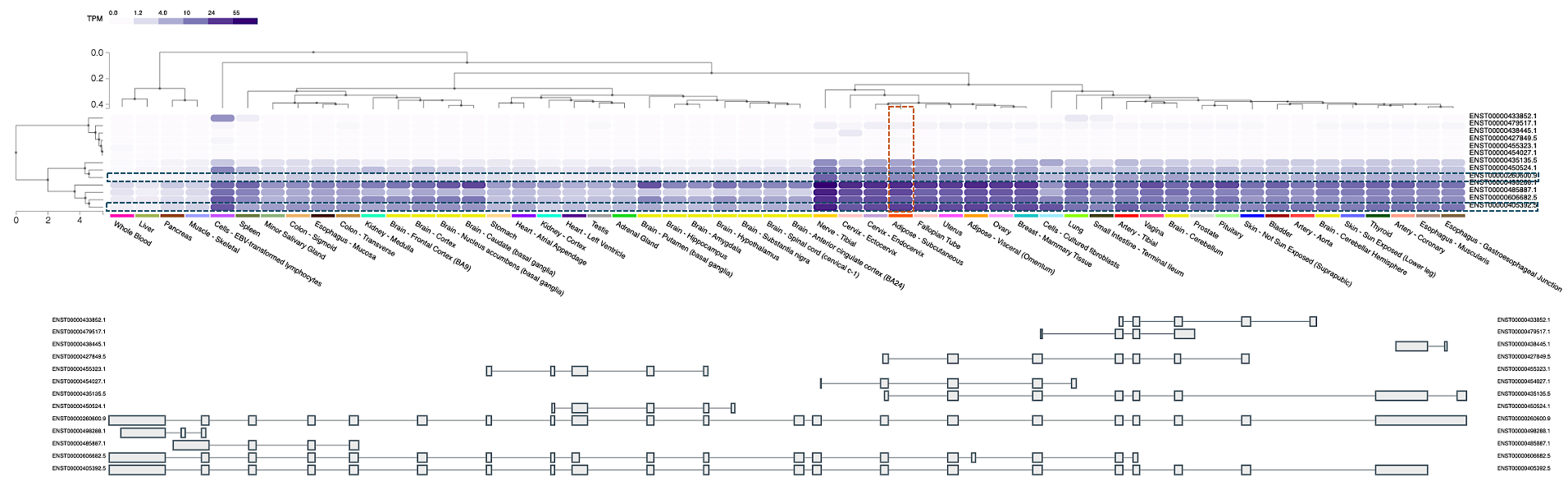

**Figure S9: Tissue-specific isoform usage of ADCY3 across human tissues.**
Heatmap showing relative expression (TPM, transcripts per million) of ADCY3 isoforms across GTEx v8 tissues. The hierarchical clustering dendrogram groups isoforms based on shared expression profiles. Adipose tissue (highlighted by the dashed box) exhibits a distinct set of ADCY3 isoforms compared to other tissues such as liver, hypothalamus, and cerebellum. These findings support a model in which tissue-specific isoform usage contributes to differential regulation of ADCY3, providing a mechanistic basis for the observed splicing QTL effects of rs11676272.

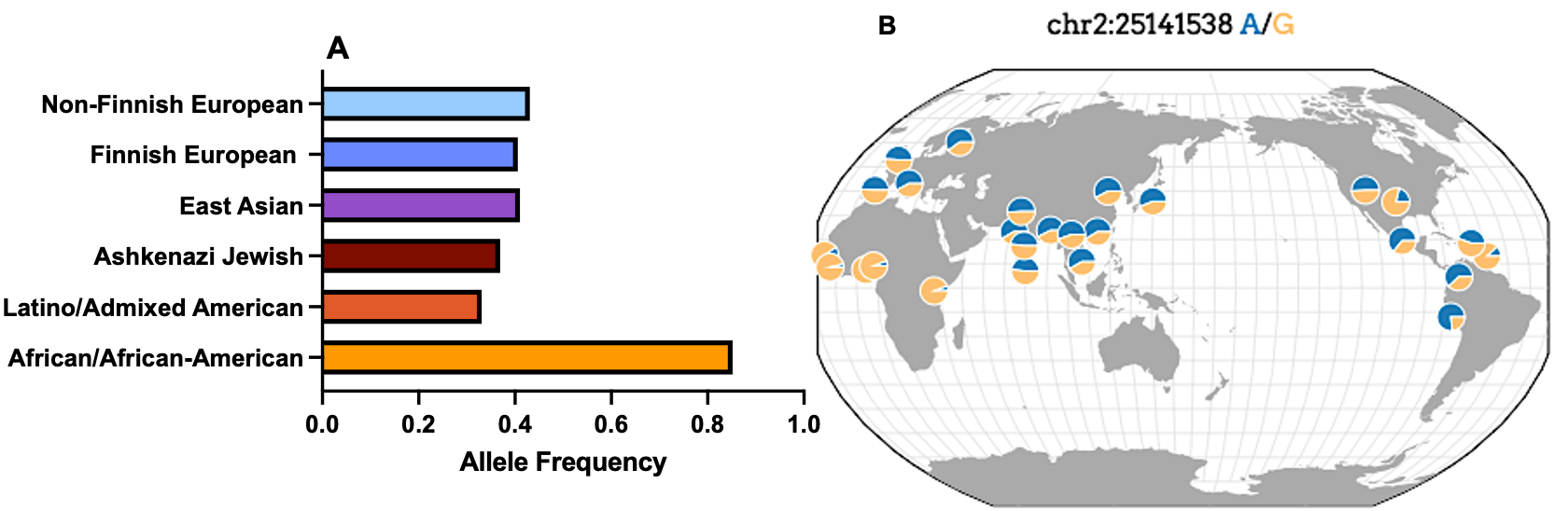

**Figure S10: Global allele frequencies and evolutionary signals for the ADCY3 rs11676272 variant.**
(**A**) Bar plot of rs11676272 allele frequencies across global populations from the gnomAD database, showing that the ancestral G (risk) allele is most common in African populations, whereas the derived A (protective) allele is more prevalent in non-African populations. (**B**) World map showing the geographic distribution of the A (blue) and G (orange) alleles in the 1000 Genomes populations. The elevated frequency of the A allele outside Africa, along with negative Tajima’s D statistics in East Asian (–1.4390) and European (–0.7118) populations, suggests a signature of recent positive selection for the protective allele, potentially reflecting adaptation to environmental pressures such as colder climates.
